# A powerful framework for differential co-expression analysis of general risk factors

**DOI:** 10.1101/2024.11.29.626006

**Authors:** Andrew J. Bass, David J. Cutler, Michael P. Epstein

## Abstract

Differential co-expression analysis (DCA) aims to identify genes in a pathway whose shared expression depends on a risk factor. While DCA provides insights into the biological activity of diseases, existing methods are limited to categorical risk factors and/or suffer from bias due to batch and variance-specific effects. We propose a new framework, Kernel-based Differential Co-expression Analysis (KDCA), that harnesses correlation patterns between genes in a pathway to detect differential co-expression arising from general (i.e., continuous, discrete, or categorical) risk factors. Using various simulated pathway architectures, we find that KDCA accounts for common sources of bias to control the type I error rate while substantially increasing the power compared to the standard eigengene approach. We then applied KDCA to The Cancer Genome Atlas thyroid data set and found several differentially co-expressed pathways by age of diagnosis and *BRAF* mutation status that were undetected by the eigengene method. Collectively, our results demonstrate that KDCA is a powerful testing framework that expands DCA applications in expression studies.

## 1 Introduction

A transcriptomic study measures the expression of thousands of genes simultaneously to help understand the molecular basis of diseases. In such studies, analyzing genes in the context of their associated pathways can reveal the underlying biological activity causing disease. In particular, a pathway that contributes to disease will co-regulate in normal conditions but, in the presence of an environmental or genetic risk factor, will result in a homeostatic breakdown. Such dysregulated pathways relate to the concept of decanalization [1, 2], where pathway coregulation evolves over many generations under stabilizing selection but is then disrupted by recent risk factors. Thus, discovering genes in a pathway whose shared expression (i.e., coexpression) differs across a risk factor can help uncover underlying biological mechanisms that induce dysregulation and cause disease susceptibility. As a result, differential co-expression analysis of pathways has proven valuable in studies of Alzheimer’s disease [3, 4], cancer [5, 6], and type 2 diabetes [7].

While identifying dysregulated pathways provides important biological insights, there are several inherent limitations to existing methods that limit their practical utility. In particular, there are methods to test for differential co-expression by assessing whether the correlation between pairs of genes in a pathway vary as a function of the risk factor (e.g., DiffCorr [8], DGCA [9], DiffCoEx [10], and GSCA [11]). However, such approaches are unable to analyze continuous risk factors (e.g., age of diagnosis, body mass index, and/or antigen levels) as well as multiple risk factors simultaneously. These methods also have difficulty accounting for batch and variance-specific effects which can increase the number of false discoveries in a study.

Lea et al. (2019) developed a method, CILP, that addresses some of these limitations but only considered a pair of genes for differential co-expression analysis rather than an entire pathway. While CILP could theoretically be applied to all pairwise combinations of genes in a pathway, it does not leverage shared information across genes and requires a multiple testing correction. These factors reduce the power to detect differentially co-expressed pathways. An alternative method that leverages shared information across genes is the standard eigengene approach [12], which tests the top principal component of an appropriately adjusted gene expression correlation matrix. However, there is no reason to assume *a priori* that the top principal component captures the desired biological signal, as it depends on many factors, including signal sparsity, pathway size, and correlation structure [13]. As such, ignoring the lower-variance components can lead to power loss and the failure to detect a signal. Collectively, these limitations have restricted the application of differential co-expression analysis in genomic studies.

We develop a flexible kernel framework to test for differential co-expression in a pathway. Our proposed method, Kernel-based Differential Co-expression Analysis (KDCA), is motivated by the observation that pathways are differentially co-expressed when the correlation patterns between genes are associated with a risk factor. KDCA harnesses such correlation patterns across all genes in a pathway while handling pathways of arbitrary size, multivariate sets of general (i.e., continuous, discrete, or categorical) risk factors, and sources of bias due to batch and variance-specific effects. Compared to the eigengene approach, KDCA does not ignore lower-variance components and leverages all information in a pathway to increase power when testing for differential co-expression.

We evaluate KDCA using comprehensive simulations that demonstrate type I error rate control under the presence of bias-inducing sources. We then show that, depending on the pathway architecture, KDCA can substantially increase power compared to the eigengene approach. Finally, we apply KDCA to The Cancer Genome Atlas (TCGA) thyroid data set [14] and discover differentially co-expressed pathways with respect to age of diagnosis and *BRAF* mutation status that were undetected by the eigengene approach. Overall, our results indicate that KDCA can flexibly test a pathway for differential co-expression using one or many risk factors while accounting for common sources of bias.

## 2 Methods

### 2.1 Overview

Differential co-expression is broadly defined as a risk factor that influences the correlation between two or more genes in a pathway. While KDCA can directly test the risk factor’s influence on pathway expression (i.e., the main effects), we focus on a more complex situation where two or more genes share an unobserved (or latent) factor that interacts with the risk factor to impact gene expression (Figure 1a; Supplementary methods). The existence of such effects stems from the complexity of gene regulation pathways where the expression levels of most genes rise and fall over time and across cells. The presence of observable factors (e.g., age or sex) can influence these fluctuations, but so can other factors that are difficult to identify and measure such as substrate abundance. The interaction between the risk factor and the unobservable factor(s) creates complex non-linear behavior that can be overlooked by only considering the main effect.

**Figure 1:**
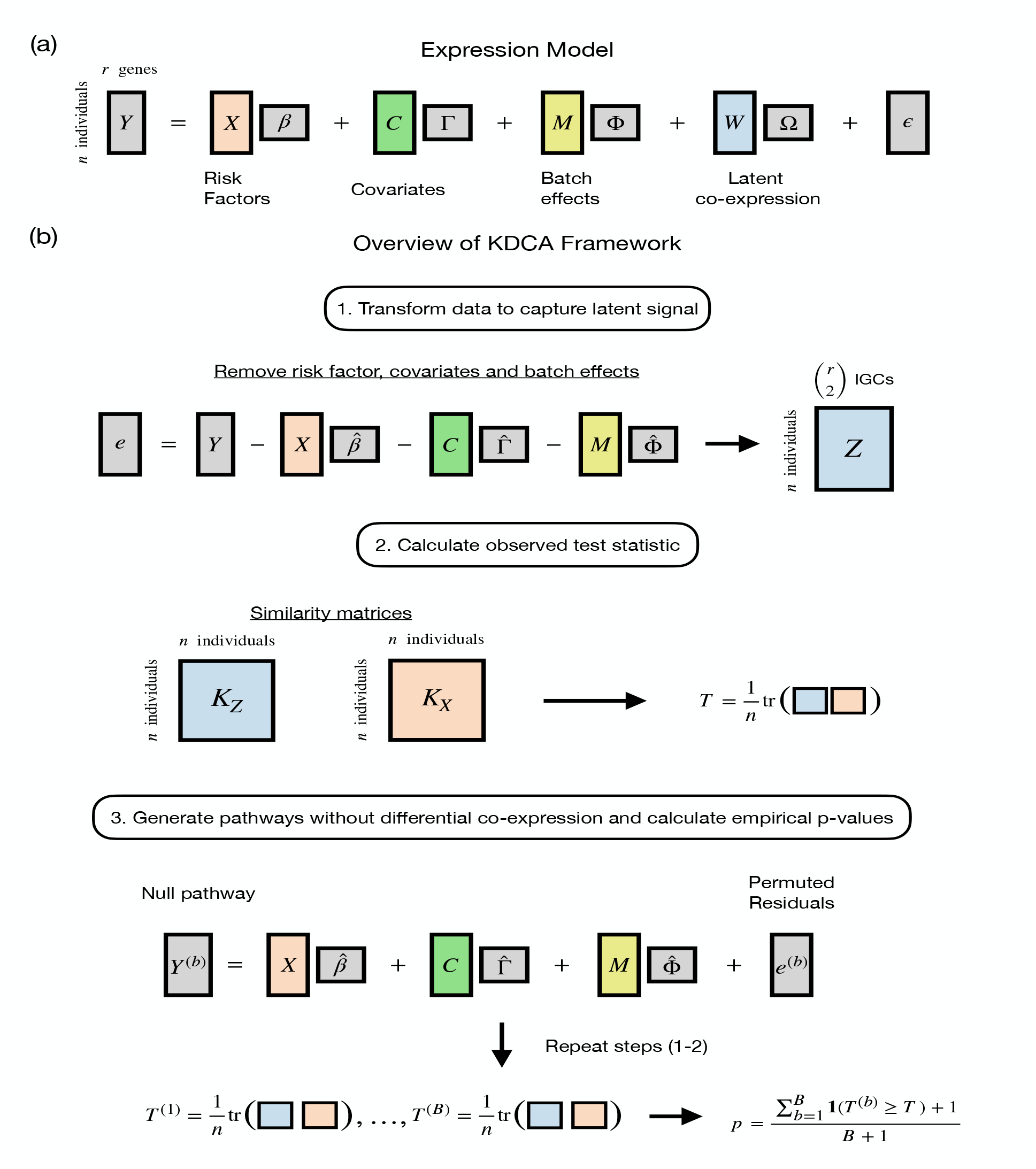
Overview of the KDCA framework. (a) The general gene expression model for a pathway with latent co-expression. (b) KDCA first calculates standardized residuals while accounting for the mean and variance effects from the risk factor, covariates and batch effects. Kernel matrices are calculated using the risk factor(s) and estimates of the individual-specific gene correlations (IGC) to construct the test statistic. A permutation algorithm is then implemented to approximate the null distribution and construct empirical *p*-values while accounting for mean and variance effects.

KDCA is based on our earlier method LIT [13], which leveraged differences in cross-trait correlation patterns to identify single-nucleotide polymorphisms with latent interactive effects in biobank-scale genome-wide association studies. In the genomics setting, KDCA instead assesses differential co-expression by relating an individual’s risk factor(s) to individual-specific gene correlations (IGC) between genes in a pathway (Figure 1b). We first estimate IGCs between pairs of genes by multiplying the residuals of different pairs of genes together, after adjusting for covariates, batch effects, and variance effects. We then test whether the elements of a matrix comprised of pairwise similarity of IGC terms is independent of the elements of a second matrix comprised of pairwise risk factor(s) similarity. The similarity between variables is measured with a user-specified kernel function, such as a linear kernel (analogous to scaled covariance), a projection kernel, or a Gaussian kernel. Since the optimal choice depends on pathway architecture (shown below), KDCA integrates multiple kernels to maximize discovery for general risk factors.

The KDCA framework differs from the LIT framework in two important ways. Firstly, LIT was developed for biobank-scale data sets rather than the modest sample sizes of gene expression data sets. Consequently, the resulting LIT test statistics rely on asymptotic theory that does not hold for these smaller data sets. In this work, we develop a novel permutation algorithm to calculate valid *p*-values for data sets of modest sample size. Secondly, LIT detected any changes in covariance from the risk factor, including from variance-specific effects. As variance-specific effects are not of interest in differential co-expression analysis, KDCA constructs the gene-by-gene correlations to avoid variance-specific false discoveries.

### 2.2 KDCA framework

Consider a pathway with expression value *Y*_*jk*_ for *j* = 1, 2, … *n* individuals and *k* = 1, 2, …, *r* genes. We assume that the expression values are influenced by a risk factor of interest, a set of measured covariates and batch effects, and a set of unobserved (or latent) factors (Figure 1a). More formally, let ***X*** = [*X*_1_, *X*_2_, …, *X*_*n*_]^*T*^ be a *n ×* 1 vector containing a general (i.e., continuous, discrete, or categorical) risk factor, ***C*** be a *n × l*_*C*_ matrix of measured covariates, ***M*** be a *n × l*_*M*_ matrix of batch effects, and ***W*** be a *n × l*_*W*_ matrix of latent factors. For simplicity, we present our algorithm with respect to a single risk factor, but the framework can easily be extended to handle multiple risk factors (see simulation results). We assume that the expression matrix, ***Y***, is approximately normally distributed: for RNA-seq data, this can be log counts per million (logCPM) or the logarithm of the expression value while adjusting for library size as a fixed effect. Our primary objective is to assess whether the risk factor ***X*** interacts with the latent factors ***W*** to influence expression in a pathway.

A risk factor that causes dysregulation in a pathway can be detected using estimates of the individual-specific gene correlations (IGC). The gene-by-gene correlation estimates for each individual (i.e., the IGC) are constructed while accounting for the mean and variance effects from the risk factor, covariates, and batch effects. Let the standardized residuals be denoted by 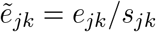, where 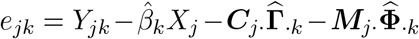 are the residuals adjusted (via least squares or double GLM) for the mean effects from the risk factor, covariates, and batch effects, and 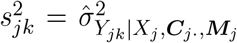 is the estimated conditional variance. Note that the subscript ‘*·*’ is a placeholder to represent all of the columns (or rows) of a matrix. The conditional variance can be estimated within each group (for a categorical risk factor) or modeled with a double GLM (for a continuous or discrete risk factor) [15]. The variance effects of other variables can also be included within the double GLM procedure. Importantly, standardizing the residuals prevents false discoveries occurring from a risk factor with only variance effects. The pathways 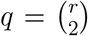 IGC terms are estimated by taking the cross products of the standardized residuals, i.e., 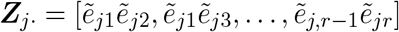.

With the cross products and risk factor, our inference goal can be summarized as follows. Under the null hypothesis of no differential co-expression, the estimates of the IGCs are independent of the risk factor:

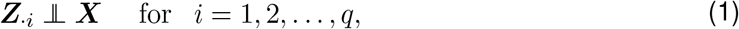

where ***Z***_·*i*_ is the cross product of the *i*th pair of genes and ‘╨’ denotes statistical independence. Intuitively, if the cross products are not independent of the risk factor, then it implies that there is some latent factor(s) that interacts with the risk factor to influence expression. While a promising strategy is to test each cross product separately [1], such an approach does not leverage shared information across tests and suffers a power loss by correcting for multiple testing.

To address these shortcomings, we implement a kernel distance covariance framework to test for differential co-expression in a pathway [16–18]. The KDCA framework uses a kernel-based independence test called the Hilbert-Schmidt independence criterion (HSIC) [16]. The HSIC generalizes well-known testing procedures in statistics (e.g., the RV coefficient [19], distance covariance [20], and multivariate distance matrix regression (MDMR) [21]) and has been used extensively in genomics [13, 22–25]. We summarize the KDCA framework for general gene expression studies below.

**Step 1: Constructing kernel matrices** The HSIC constructs a test statistic that measures the amount of shared signal between a similarity matrix for the risk factor and a similarity matrix for the cross products. To calculate a similarity matrix, a kernel function is specified to measure the similarity between the *j*th and *j*′th individual. There are many options for the kernel function, such as the linear kernel (equivalent to a scaled covariance) 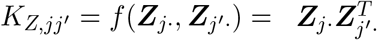, the projection kernel 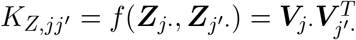, where ***V*** is the left singular vectors of ***Z***, and the Gaussian kernel 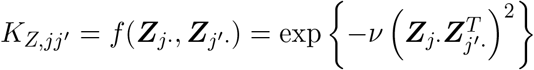, where *ν* is a tuning parameter [25,26]. The optimal kernel depends on the structure (or architecture) of the pathway. For example, the Gaussian kernel is suitable for complex non-linear co-expression patterns while the linear and projection kernels are more suitable for linear co-expression patterns. However, even when linear relationships are expected, the choice between the linear and projection kernels is not clear (see ref. [13] for a thorough investigation). Since the optimal kernel choice is unknown *a priori* in our setting, we evaluate the linear kernel, the projection kernel, and the Gaussian kernel for the cross products. For the risk factor, we only consider the linear kernel *K*_*X,jj′*_ = *f* (***X***_*j*_, ***X***_*j′*_) = *X*_*j*_*X*_*j′*_. We denote the similarity matrix of the cross products and risk factor as ***K***_*Z*_ and ***K***_*X*_, respectively. In the Gaussian kernel implementation, we set *ν* = 0.0001 as it performed well across all simulation settings.

**Step 2: Test statistic and inference** With the similarity matrices ***K***_*Z*_ and ***K***_*X*_, the HSIC test statistic measures the ‘overlap’ between the two matices as

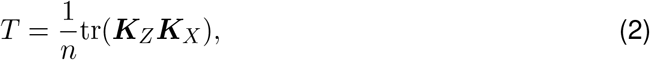

where small values of *T* suggest independent matrices (no differential co-expression) and large values of *T* suggest that the pairwise elements of the matrices are dependent (differential co-expression). Under the null hypothesis, the HSIC test statistic follows a weighted sum of Chi-squared random variables, i.e., 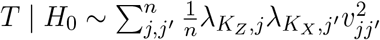, where 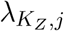 and 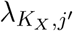 are the ordered non-zero eigenvalues of the respective matrices and *v*_*jj′*_ ∼ Normal(0, 1). For biobanks or other studies with large sample sizes, we can approximate the null distribution in a computationally efficient manner using Davies’ method [24, 25, 27]. However, there are not many expression studies that have sample sizes sufficiently large for this approximation to hold. Although there exists a small sample size approximation for a related test statistic that estimates the null distribution by matching the moments of the permutation statistics to a Pearson type III distribution [28], the underlying assumptions do not hold in our setting since we remove the mean and variance effects of the risk factor from the gene expression values (shown in simulations).

We instead develop a permutation algorithm to estimate the empirical null distribution of the HSIC test statistic. For *b* = 1, 2, …, *B* permutations, let ***H***^(*b*)^ denote a *n × n* matrix that permutes the rows of the standardized residuals, i.e., 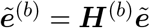 is a permuted set of residuals that removes any differential co-expression but conserves the gene-by-gene correlations. Intuitively, since the permutation shuffles the expression values for each gene in the same order (row-wise), the correlation between genes does not change. Instead, the row-wise permutation removes any associations between the risk factor and the latent factors (i.e., co-expression) to generate a pathway without differential co-expression. Our permutation algorithm is summarized as follows.

1. For *b* = 1, 2, …, *B* permutations, generate the pathway under the null hypothesis:

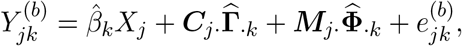

Where 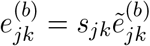.

2. Calculate the cross products and construct the corresponding kernel matrix 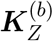. The null statistic is calculated as 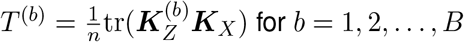.

3. Using the observed statistic *T* and null statistic *T* ^(*b*)^, calculate the empirical *p*-value according to

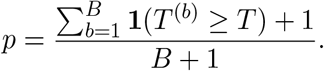

An important feature of the above algorithm is that the mean effects from the risk factor, covariates, and batch effects are preserved when generating the null pathway. Furthermore, we account for the influence of the risk factor on expression variance to avoid false discoveries (i.e., not differentially co-expressed) from factors with strong variance effects. While we emphasize a single risk factor here, the algorithm can be extended to incorporate multiple risk factors (whether continuous, discrete, categorical, or a mixture of all three), and also adjust for variance-specific effects from such factors.

### 2.3 Combining information across kernels to maximize power

KDCA requires choosing a kernel function for the cross products and risk factor. However, it is unclear which kernel is optimal *a priori* given that they capture different measures of coexpression similarity. Since one kernel may outperform the others, we consider an aggregate implementation to maximize the statistical power and adjust the above algorithm as follows.

We first calculate the observed test statistic using the linear, projection and Gaussian kernels (although other kernels could be added). We then implement the permutation algorithm such that the null statistics and corresponding empirical *p*-values are calculated separately for each kernel. Calculating the empirical *p*-values before combining the kernels standardizes the null distributions of each kernel. We then construct a test statistic that combines the empirical *p*-values using Fisher’s method, i.e., 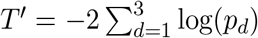, where *p*_*d*_ is the empirical *p*-value for the *d*th kernel. To estimate the null distribution of this statistic, we combine the empirical *p*-values in permutation *b* as 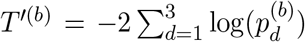. Finally, an empirical *p*-value for the observed statistic *T* ′ is calculated using the combined null statistics *T* ′^(*b*)^. It is important to note that any *p*-value combination approach can be used, although we found Fisher’s method performed well in practice.

### 2.4 Simulation study

We evaluated the performance of KDCA using simulated data. We considered a categorical and continuous risk factor: the categorical variable was generated as *X*_*j*_ ∼ Bernoulli(0.5) and the continuous variable was generated as *X*_*j*_ ∼ Uniform(0, 2). For the pathway, we assumed the risk factor had the following effect on the mean and variance of each gene, and on the covariance between each pair of genes:

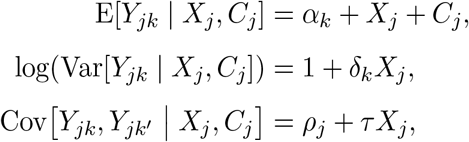

where *α*_*k*_ ∼ Normal(0, 1) is the intercept value; *C*_*j*_ ∼ Uniform(0, 2) is an observed covariate; *ρ*_*j*_ ∼ Uniform(0.25, 0.50) is the baseline correlation between each pair of genes; *δ*_*k*_ ∼ Uniform(0.0, 0.1) is the variance-specific effect size of the risk factor; and *τ* ∼ Uniform(0.1, 0.2) is the correlation-specific effect size of the risk factor.

We considered various configurations of differential co-expression due to the risk factor. We first set the sample size to be *n* = 300 and pathway size to be *r* = 10 or 50 genes. Note that the average pathway size of the BioCarta pathway database [29] in our applied example is 8.3, and so *r* = 50 reflects larger pathways. We then varied the proportion of genes in a pathway with no differential co-expression as 0, 0.20, 0.40, 0.60, and 0.80. This represents different sparsity levels of differential co-expression. We also considered a more complex pathway correlation pattern where the direction of the effect sizes was randomly assigned to be positive or negative (i.e., *±τ*). In total, we simulated 200 replicates at each configuration and the empirical power was calculated at a significance threshold of 0.05.

To evaluate type I error rate control under the null hypothesis, we considered two different settings. The first setting evaluated the null in the above simulations by assigning *τ* = 0 for all pathways. The second setting simulated 10,000 genes from a negative binomial distribution with the following parameters:

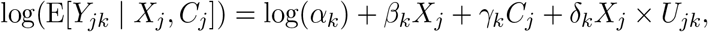

where log(*α*_*k*_) ∼ Lognormal(5.54, 0.697) is the baseline count size for the *k*th gene; *β*_*k*_ ∼ Normal(0, 0.01) is the effect size of the risk factor; *γ*_*k*_ ∼ Normal(0, 0.01) is the effect size of the covariate; and the latent interaction *X*_*j*_ *× U*_*jk*_ represents a variance-specific effect where *U*_*jk*_ ∼ Normal(0, 1) with effect size *δ*_*k*_ ∼ Normal(0, 6.25 *×* 10^−4^). We scaled the counts to reflect varying library sizes where the total counts for each sample was randomly chosen to be 2 *×* 10^6^ or 10 *×* 10^6^. The dispersion parameter of the negative binomial distribution was assigned to be 0.25 for each gene in order to reflect human data sets. We applied a log2 transformation to the counts and treated the estimated (log-transformed) library size as a fixed covariate. In both settings, we simulated 20 data sets with 1,000 pathways of size *r* = 10 or 50. The type I error rate of KDCA was evaluated at a threshold of 0.05 with *B* = 1,000 permutations.

To assess the performance of KDCA with multiple risk factors, we simulated risk factors as *X*_*jl*_ ∼ Bernoulli(0.5) for *l* = 1, 2, 3 factors. The expression values were generated by incorporating the risk factors into the above model as 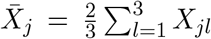. We then applied KDCA to the multiple risk factors to assess the performance under a multivariate setting.

### 2.5 TCGA study

The Cancer Genome Atlas (TCGA) is a large-scale genomic program to characterize various cancer types in humans. For our analysis, we used the RNA-seq data from thyroid carcinoma (THCA) samples downloaded from recount3 [30]. We tested for evidence of differential co-expression using *BRAF* mutation status (extracted from TCGAbiolinks [31]) and age of diagnosis. The *BRAF* mutation is a prognostic marker for papillary thyroid cancer (PTC) and associated with overall worse prognosis [32], while a patient’s age of diagnosis is a known risk factor for many cancers as mortality increases with age [33, 34]. We restricted our analysis to primary tumor samples and removed any formalin-fixed paraffin-embedded (FFPE) samples. We then removed non-protein coding genes and any genes with log2 counts per million (logCPM) values less than 1 in 50% of the samples. We also ensured that there were no over-lapping genes in the data set. After filtering, we selected 5,000 genes with the largest variation (logCPM) across 494 samples.

Following previous recommendations [35], we estimated latent batch effects using the log2-transformed counts and found 28 latent variables in our data set (treated as adjustment variables). This is a conservative approach to differential co-expression testing as it may remove the biological signal of interest. However, it was previously shown to outperform models that explicitly modeled sources of confounding (e.g., RIN, exonic rate, or GC bias) in gene network reconstruction [35], suggesting that there are diverse sources of confounding in expression data that are challenging to account for. We then used the canonical BioCarta pathways [29] from the Molecular Signatures Database [36] where pathway sizes less than 5 were removed. In total, there were 162 pathways with an average tested set size is 8.3 (Figure S1). Finally, we implemented KDCA and the eigengene approach using 10,000 permutations.

## 3 Results

### 3.1 KDCA controls the type I error rate

We performed comprehensive simulations to assess the type I error rate of KDCA (Section 2.4). In particular, we simulated pathways of size 10 and 50 with a sample size of 300 under two settings. The first setting generated pathways assuming the data followed a multivariate normal data while the second setting generated pathways assuming the data followed a negative binomial distribution. In the latter setting, we applied a log2 transformation to the counts and treated library size as a fixed effect. In both cases, the continuous and categorical risk factor impacted the mean and variance of the expression values.

We find that our permutation algorithm provides type I error rate control in either the normal or negative binomial simulation cases, regardless of the kernel choice or risk factor type (Figure S2, S3). Additionally, our implementation that combines information across the different kernels also provided type I error rate control. Finally, when there are multiple risk factors included in the model, we find that all of the KDCA implementations control the type I error rate (Figure S4). Thus, our permutation approach, which models the mean and variance effects from the risk factor, controls the type I error rate in the presence of variance-specific effects for continuous or categorical risk factors.

We also evaluated an approximation of the theoretical null distribution for a related test statistic [28] in the normal setting. For categorical risk factors, we find that the type I error rate is very conservative for small and large pathway sizes (Figure S5). Alternatively, for continuous risk factors, the type I error rate is not controlled. Note that the mean and variance effects of the risk factor are adjusted from the genes before kernel construction. This violates a key assumption of the approximation, i.e., permuting one kernel does not impact the other kernel under the null. While there is a large sample size approximation to the KDCA test statistic [27], most expression studies do not have large enough sample sizes for this approximation to hold. As such, we find that our permutation-based approach for calculating valid *p*-values is required in this setting, even though it increases the computational time (see Figure S6 for computational time comparisons).

### 3.2 The optimal kernel choice depends on pathway architecture

We assessed the power of KDCA in our simulation study using various kernel choices. We varied the proportion of genes that were differentially co-expressed in a pathway in the categorical and continuous risk factor settings. We considered a ‘Positive’ scenario where the effect size direction of the latent co-expression was the same and a ‘Positive/Negative’ scenario where the direction was randomly positive or negative. We applied KDCA using a linear kernel, a Gaussian kernel, a projection kernel, and an implementation that aggregates the statistics using Fisher’s method. Importantly, the first three KDCA implementations are different measures to capture the signal while the last attempts to maximize the power by leveraging information across the kernels.

Our simulation results suggest that the best kernel choice depends on the pathway size, sparsity of the signal, and correlation structure (i.e., ‘Positive’ or ‘Positive/Negative’; Figure 2). In particular, for a continuous risk factor, we find that the Gaussian and linear kernels outperform the projection kernel (Figure 2a). However, for a categorical risk factor, the projection kernel outperforms the Gaussian and linear kernel in the ‘Positive/Negative’ setting with a pathway size of 10 (Figure 2b).

**Figure 2:**
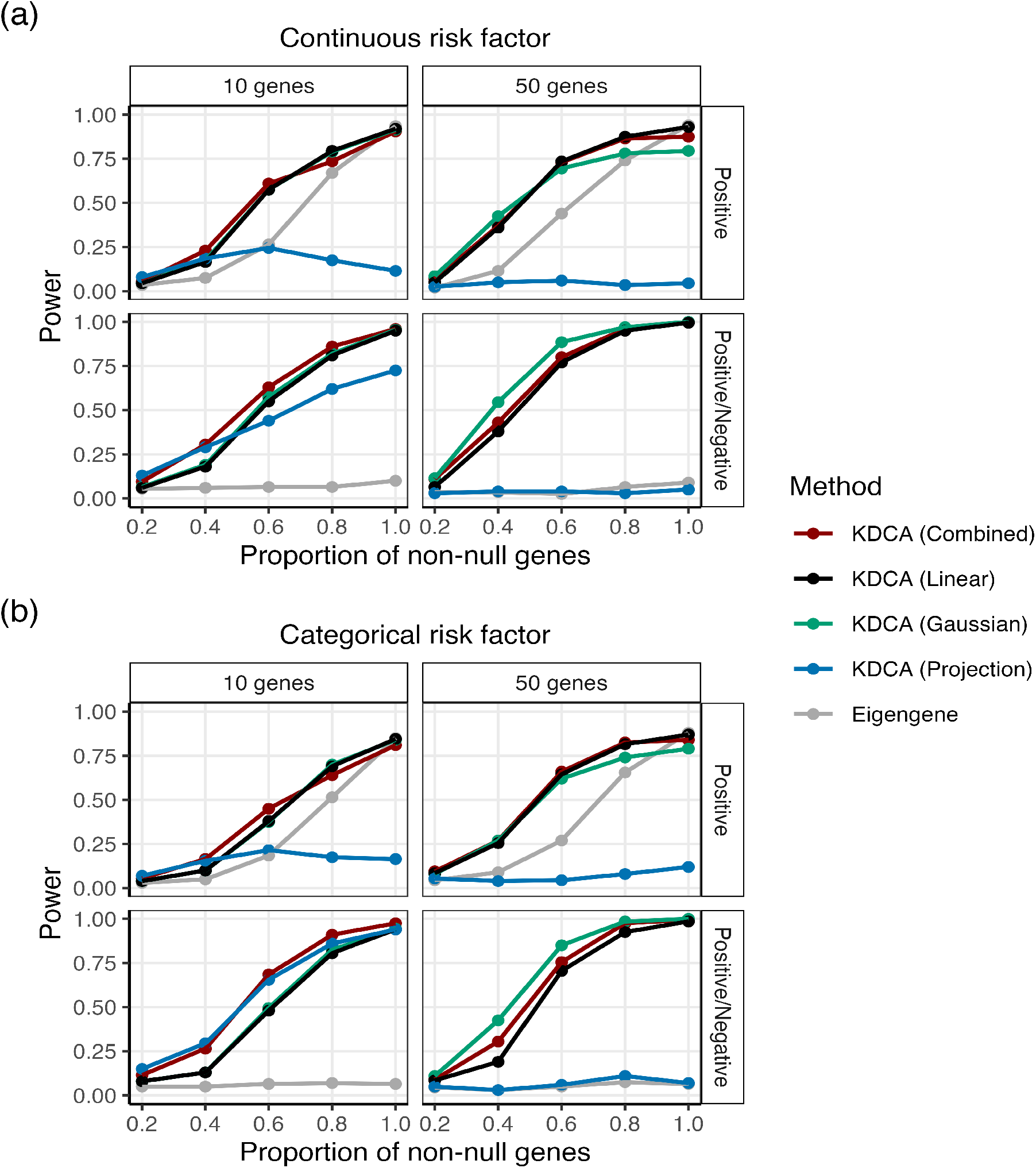
The empirical power of KDCA using the linear (black), Gaussian (green), and projection (blue) kernels when the primary variable is (a) continuous and (b) categorical. We also compared an implementation of KDCA that maximized power across all kernels (combined; red) to a standard eigengene (grey) approach. Our simulation study varied the pathway size (columns) and the type of differential co-expression (rows). Each point is the empirical power from 200 simulations at a significance threshold of 0.05.

Since the best performing kernel is unclear, we implemented the KDCA combined approach that aggregates information across the three kernels to maximize statistical power. We find that KDCA combined provides the best performance for maximizing power across the various settings. We also found similar results when considering multiple risk factors (Figure S7).

### 3.3 KDCA improves power compared to the eigengene approach

We compared KDCA to the standard eigengene approach that tests the top eigenvector of the cross product matrix (adjusting for the mean and variance effects of the primary variable). To construct an empirical *p*-value, we apply the same permutation algorithm as KDCA. In general, the linear kernel outperforms the eigengene approach because it aggregates information from all pairwise correlations of genes within a pathway (Figure 2, S7). Similarly, the Gaussian kernel tends to outperform the eigengene approach. Finally, while the projection kernel outperforms the eigengene method in the ‘Positive/Negative’ setting, it underperforms in the ‘Positive’ setting.

Our findings can be interpreted by considering the singular value decomposition of the cross product linear kernel matrix. In particular, the eigengene approach is equivalent to testing the top eigenvector of the linear kernel matrix. Therefore, when the eigengene approach performs comparably to KDCA with a linear kernel, it suggests that the signal of interest is captured by the top eigenvector. Alternatively, when the linear kernel outperforms the eigengene approach, it suggests that the signal is captured at the lower eigenvectors. For example, in the ‘Positive/Negative’ setting where the signal is captured by lower eigenvectors, KDCA with a linear kernel can substantially outperform the eigengene approach. While the power of the eigengene approach improves in the ‘Positive’ setting where the signal is expected to be captured by the top eigenvector, it is still lower than KDCA with a linear kernel. This suggests that the signal is also captured by the lower eigenvectors. Note that, when the signal is only captured by a small number of eigenvectors, the projection kernel can have the lowest power because it treats the eigenvectors in the linear kernel matrix equally.

### 3.4 KDCA reveals differentially co-expressed pathways in TCGA analysis

Thyroid cancer is a common cancer with an incidence rate that has been increasing over the past few decades [37]. While the overall survival rate is very promising with early treatment, the cancer is heterogeneous and certain subtypes can be lethal [14]. To help elucidate biological variation not captured by traditional differential expression analysis, we applied the KDCA framework to The Cancer Genome Atlas (TCGA) thyroid cancer data set. Our analysis focused on papillary thyroid carcinomas (PTCs) samples with two risk factors: *BRAF* mutation status (a categorical risk factor) and age of diagnosis (a continuous risk factor).

We find that KDCA combined is substantially more powerful than the eigengene approach, in line with our simulation study (Figure 3; Table S1). In particular, at a FDR threshold of 0.15, the eigengene approach does not detect any differentially co-expressed pathways while KDCA combined detects 3 and 6 pathways for *BRAF* mutation status and age of diagnosis, respectively. The individual kernel implementations also outperform the eigengene approach (Figure S8), demonstrating that KDCA benefits from aggregating all the information from the cross product matrix.

**Figure 3:**
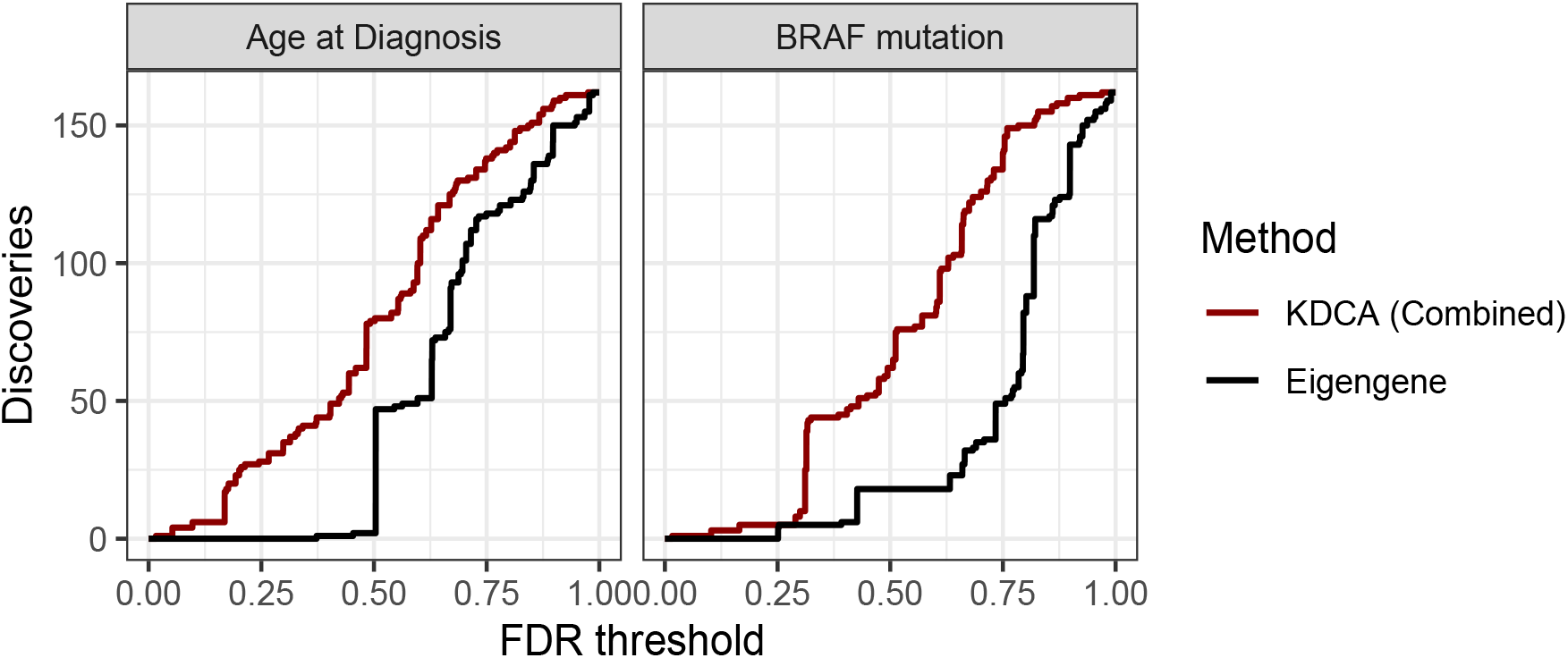
The number of detections as a function of FDR threshold using the BioCarta pathway database in the TCGA thyroid cancer data set. We compared an aggregate version of KDCA (red) to a standard eigengene approach (grey) when the primary variable was age of diagnosis (left) and *BRAF* mutation status (right). There were a total of 162 pathways tested for differential coexpression.

When focusing on the discoveries from KDCA combined at a FDR threshold of 0.15 (Table 1), we find that the differentially co-expressed pathways are involved in cellular proliferation, the immune microenvironment, and/or are known prognostic markers. In particular, when the risk factor is age of diagnosis, the IRL2, RAS, and TFF2 pathways are differentially coexpressed. The IRL2 pathway is a set of genes involved in immune-related signaling for cellular activation and growth. The RAS signaling pathway is involved in the immune microenvironment and the *RAS* mutation is a prognostic marker [38]. Finally, the TFF2 pathway promotes cellular proliferation and can be a key component in the MAPK signaling pathway [39]. Importantly, immune-related pathways are expected as inflammation is a known hallmark of aging and can have tumor promoting effects [33].

**Table 1:**
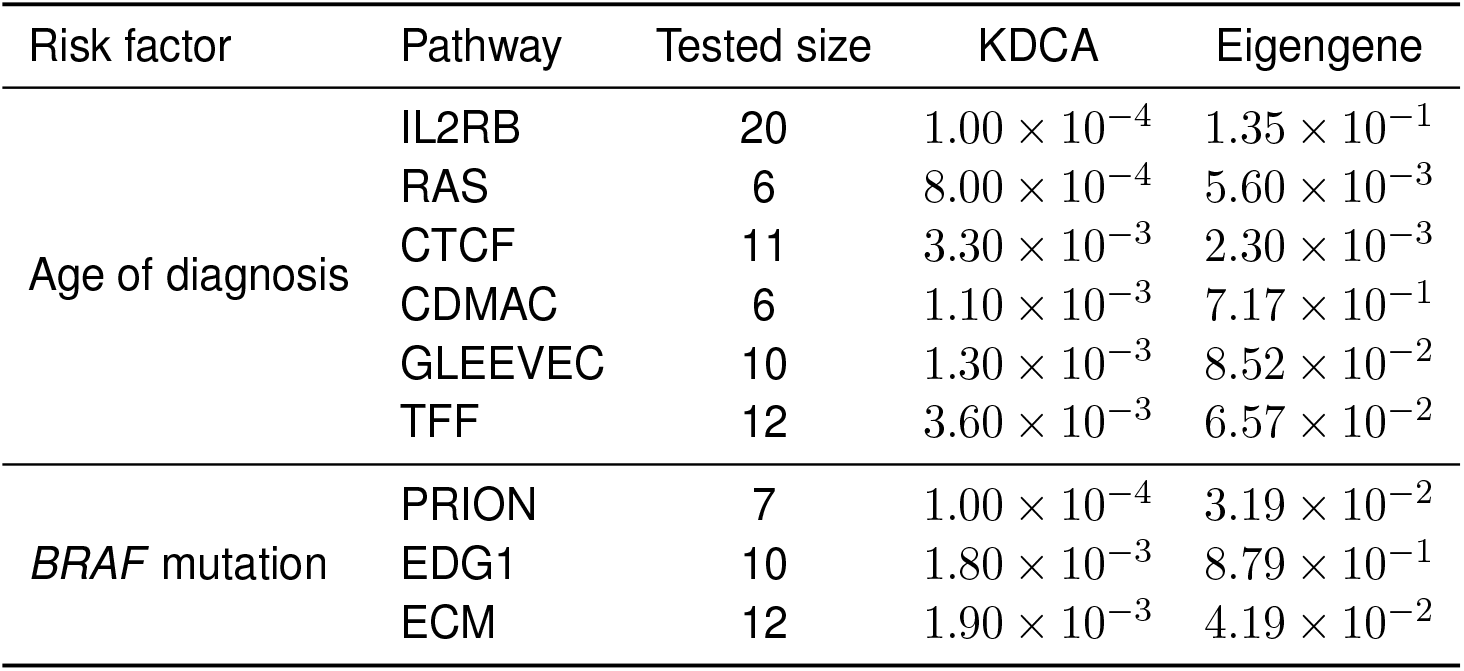
Differential co-expression analysis of the TCGA thyroid cancer data using age of diagnosis and *BRAF* mutation as risk factors of interest. KDCA combined *p*-values are reported for pathways below a FDR threshold of 0.15. The standard eigengene approach did not find any pathways below the significance threshold.

| Risk factor | Pathway | Tested size | KDCA | Eigengene |
| --- | --- | --- | --- | --- |
| Age of diagnosis | IL2RB | 20 | $1.00 \times 10^{-4}$ | $1.35 \times 10^{-1}$ |
| | RAS | 6 | $8.00 \times 10^{-4}$ | $5.60 \times 10^{-3}$ |
| | CTCF | 11 | $3.30 \times 10^{-3}$ | $2.30 \times 10^{-3}$ |
| | CDMAC | 6 | $1.10 \times 10^{-3}$ | $7.17 \times 10^{-1}$ |
| | GLEEVEC | 10 | $1.30 \times 10^{-3}$ | $8.52 \times 10^{-2}$ |
| | TFF | 12 | $3.60 \times 10^{-3}$ | $6.57 \times 10^{-2}$ |
| *BRAF* mutation | PRION | 7 | $1.00 \times 10^{-4}$ | $3.19 \times 10^{-2}$ |
| | EDG1 | 10 | $1.80 \times 10^{-3}$ | $8.79 \times 10^{-1}$ |
| | ECM | 12 | $1.90 \times 10^{-3}$ | $4.19 \times 10^{-2}$ |

When the risk factor is *BRAF* mutation status, we find EDG1, PRION, and ECM pathways are differentially co-expressed. These pathways are known to affect cell growth and survival in cancer [40–42]. In addition, we find that *MAPK1, PIK3CA, PIK3CG*, and *PIK3R1* are the top four most occurring genes when the risk factor is *BRAF* mutation status, while for age of diagnosis the top four are *HRAS, PIK3CA, PIK3CG*, and *PIK3R1*. The *MAPK1, HRAS*, and *PIK3CA* genes are associated with distinct thyroid cancer subtypes [43], suggesting the intriguing hypothesis that the unobserved latent factors are germline or somatic changes in these genes themselves. Overall, we find distinct co-expression patterns among pathways with respect to *BRAF* mutation status and age of diagnosis, indicating that the cancer samples are functionally heterogeneous.

## 4 Discussion

Our proposed kernel-based framework, KDCA, harnesses correlation patterns in a pathway to test for differential co-expression. KDCA can be applied to one or many general risk factors while accounting for covariates, batch effects, and variance-specific effects. We found that the optimal kernel choice for KDCA depends on the pathway size, correlation structure, and sparsity level. This demonstrates the complexity of detecting differential co-expression and agrees with our previous work which found the optimal kernel can also vary based on other features, such as sample size, signal-to-noise ratio at each principal component, and baseline correlation between genes (see ref. [13] for a thorough investigation). Since these factors are unknown *a priori*, we developed an implementation to maximize the power by aggregating information across kernels.

One important feature of KDCA is that it helps prevent false discoveries from variance-specific effects of the risk factor. In particular, our permutation algorithm models the mean and variance effects for general risk factors to control the type I error rate. While KDCA can handle continuous and discrete risk factors, it requires careful specification of how the factor influences the variance in the double GLM. Otherwise, it is possible that variance-specific effects from the risk factor can lead to false discoveries. More generally, we caution against rank-based transformations of expression data as these can induce variance and covariance-specific effects and lead to false discoveries.

There are a few of important observations to consider when applying the KDCA framework. First, pathways should be carefully selected in order to avoid a multiple testing problem. In this work, we focused on the BioCarta pathway database and a subset of the 5,000 most varying genes. Second, unmeasured batch effects impact the covariance between genes in a pathway, leading to false discoveries when such effects are associated with the risk factor. While a method has been proposed to capture batch effects for specific types of pathway networks [35], such an approach can also remove the biological signal of interest. In general, accounting for latent confounding variables is a very difficult problem for differential co-expression methods. As such, even though KDCA provides a framework to incorporate batch effects, false discoveries due to latent non-biological sources of variation cannot be ruled out. Finally, the computational time of KDCA increases as the pathway size and/or sample size increases (Figure S6). If running a large number of pathways with a sample sizes much larger than considered here (≫ 300), we recommend distributing pathways across multiple cores to increase computational efficiency. Notably, for biobank-scale data sets, the closed-form approximation of the null distribution can be used instead of the permutation algorithm to substantially decrease the computational time [24, 25, 27].

Differential co-expression analysis provides biological insights into expression dysregulation of a pathway. However, there are many methodological challenges that have limited the utility of such approaches, such as testing continuous or discrete risk factors, avoiding false discoveries from variance-specific effects, and modeling batch effects. We developed the KDCA framework to help address these limitations, thereby providing a powerful framework to identify differentially co-expressed pathways in expression studies.

## Software and data

KDCA is publicly available in the R package kdca. The package can be downloaded at https://github.com/ajbass/kdca. The code to reproduce the results in this work can be found at https://github.com/ajbass/kdca_manuscript and the RNA-seq data from the THCA samples can be downloaded from the recount3 package [30].

## Acknowledgements

This work was supported by NIH grants R01 AG071170 (AJB, DJC, MPE). The applied results shown here are based on data generated by the TCGA Research Network: https://www.cancer.gov/tcga.

## 5 Supplementary materials

### 5.1 Supplementary methods

#### 5.1.1 A general model for differential co-expression analysis

Differential co-expression in a pathway is a statistical phenomenon that arises from unmodeled biological variation involving an observed risk factor of interest and some unobserved (or latent) factor. Consequently, this unmodeled variation induces a correlation pattern between pairs of genes in a pathway that varies as a function of the risk factor. We first motivate the observation that such correlation patterns are due to a latent factor interacting with a risk factor, and then extend our discussion to a more general model.

##### Latent factors can induce differential correlation patterns between two genes

To simplify our discussion, consider the expression of two genes in a pathway denoted by *Y*_*j*1_ and *Y*_*j*2_ for *j* = 1, 2, … *n* individuals. Let the risk factor of interest be denoted by *X*_*j*_ and the shared latent factor be denoted by *W*_*j*_. The risk factor can be a continuous (e.g., age) or categorical (e.g., SNP or biological condition) variable, while the shared latent factor is some unobserved environmental factor that is difficult to measure. We assume that the risk and latent factors contribute to expression additively as

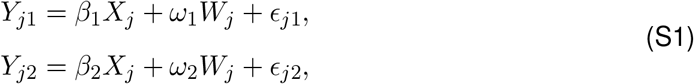

where *β*_1_ and *β*_2_ are the effect sizes of the risk factor, *ω*_1_ and *ω*_2_ are the effect sizes for the latent factor, and *ϵ*_*j*1_ and *ϵ*_*j*2_ are independent normally distributed random errors for gene 1 and 2, respectively.

If the latent factor interacts with the risk factor, and affects the expression of both genes, then a correlation pattern between the genes is induced. More specifically, assuming that the random errors are independent of the risk and latent factors, the individual-specific gene correlation (IGC) is

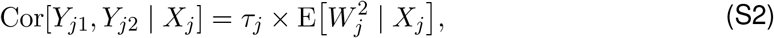

Where 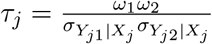 is the effect size and 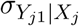 and 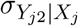 are the conditional standard deviations of gene 1 and 2, respectively. As an example, suppose the latent factor *W*_*j*_ = *X*_*j*_*U*_*j*_ is an interaction between the risk factor and an environmental factor, *U*_*j*_, that follows a standard normal distribution. In this case, when the two genes share the latent factor (i.e., *ω*_1_ ≠ 0 and *ω*_2_ ≠ 0), the conditional correlation pattern 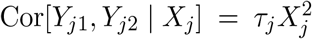 is induced and the pathway is classified as “differentially co-expressed.” Note that the conditional variance is 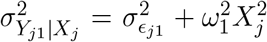 and 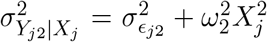 for gene 1 and 2, respectively. Thus, our strategy to assess differential co-expression is to test whether the IGCs varies as a function of the risk factor in a pathway.

##### Inferring differential correlation patterns in a pathway

We extend the previous model to include multiple genes in a pathway. Consider a pathway with expression value *Y*_*jk*_ for *j* = 1, 2, … *n* individuals and *k* = 1, 2, …, *r* genes. Let ***C*** be a *n × l*_*C*_ matrix of covariates and ***M*** be a *n × l*_*M*_ matrix of batch effects. The gene expression model is

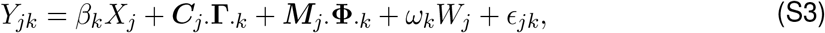

where **Γ**_·*k*_ and **Φ**_·*k*_ are effect sizes for ***C***_*j*·_ and ***M***_*j*·_, respectively, *ϵ*_*jk*_ is a normally distributed random variable whose variance can depend on the risk factor (i.e., heteroskedasticity), and the subscript ‘·’ is a placeholder to represent all of the columns (or rows) of a matrix. To illustrate our strategy, we assume the effect sizes are known. We can then calculate the residuals as

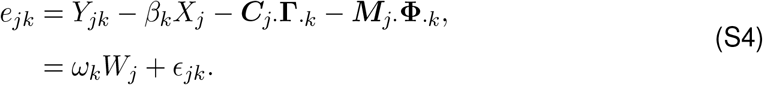

Since the risk factor may induce variance-specific effects, the residuals should be standardized to prevent false discoveries for differential co-expression testing, i.e., 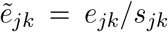 where 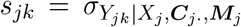*·* is the conditional standard deviation. Note that variance effects from other terms should also be modeled when standardizing the residuals.

We show in the main text that a latent interaction can be detected based on calculating the individual-specific gene covariances (IGC). For the *j*th individual, the conditional *r × r* individual-specific pathway covariance matrix is

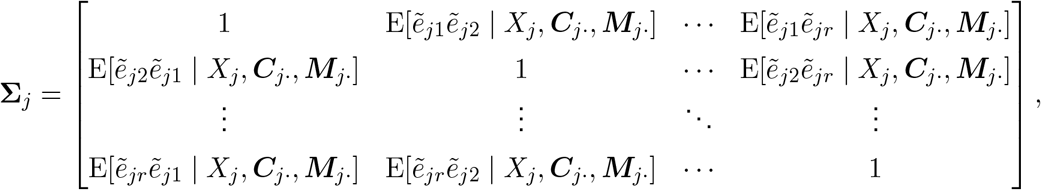

where the IGC terms are the 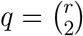 off-diagonal elements. Using the results from the previous section, the IGC between the *k*th and *k*′th genes is

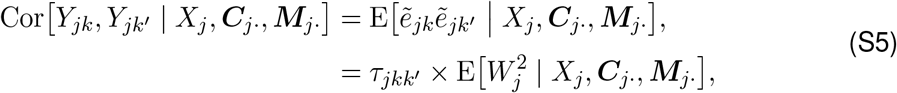

where 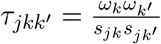 is the effect size and *s*_*jk*_ and *s*_*jk′*_ are the conditional standard deviations of gene *k* and *k*′, respectively. A pathway is differentially co-expressed if *τ*_*jkk′*_ ≠ 0 for at least one IGC term. Our results imply that the latent factor *W*_*j*_ can induce correlation patterns in a pathway whenever the factor is correlated with the risk factor and impacts the expression of the genes. Note that the above results can easily be extended to include multiple general risk factors and latent factors.

The KDCA framework estimates the IGC terms in a pathway by taking the cross products of the standardized residuals. We then use the cross products (see ref. [13] for theoretical details) to test for any evidence of differential co-expression in at least one of the IGC terms. One important topic when using KDCA involves the latent factor *W*_*j*_. The latent factor(s) is a key component in any differential co-expression analysis as it captures unmeasured variability that can impact the correlation between genes. A challenge is that a latent factor(s) can represent biological and/or technical (i.e., batch) variability. Ideally, the technical factors are included in the model as an adjustment variable to avoid false discoveries. However, distinguishing latent technical and biological factors is a very difficult problem, and it is currently unclear how to appropriately account for technical factors (without removing biological variation) in differential co-expression analysis. Here, we inferred the technical factors using the approach in ref. [35] and included them as adjustment variables in our procedure. While this approach helps control for false discoveries due to technical factors, it may also remove some of the biological factors.

### 5.2 Supplementary figures

**Figure S1:**
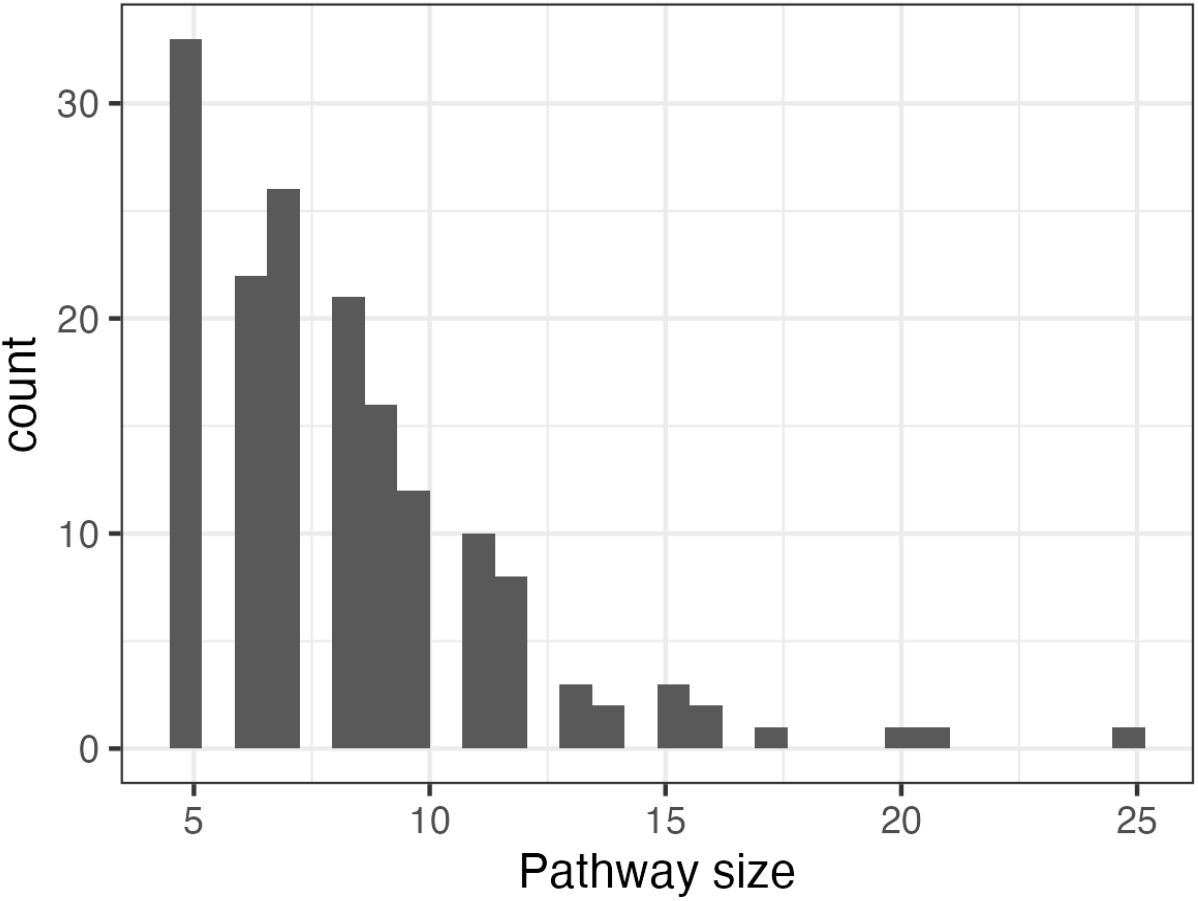
A histogram of the pathway sizes (i.e., number of genes) from the BioCarta database.

**Figure S2:**
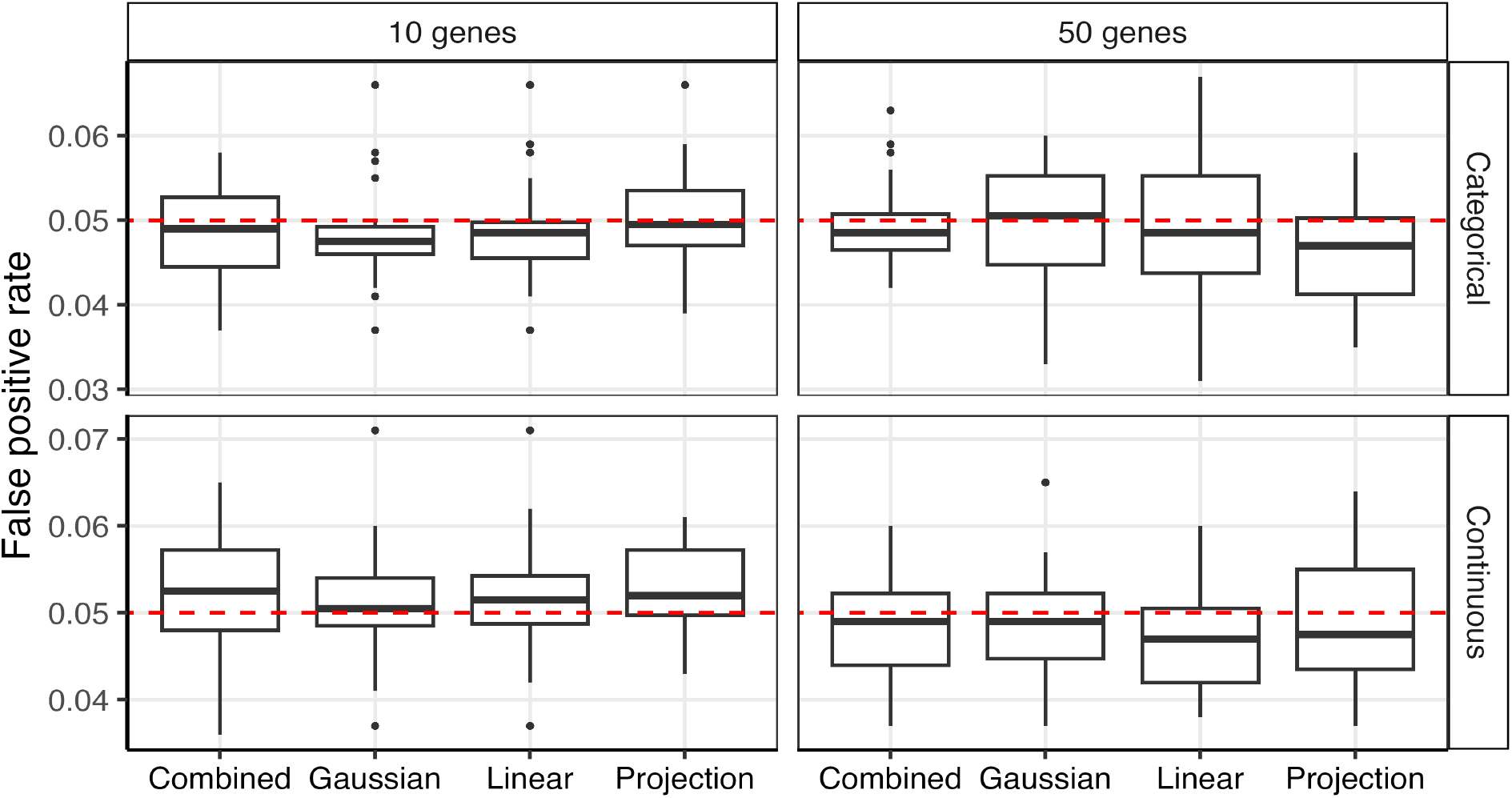
False positive rate of KDCA under the null hypothesis of no differential co-expression using the Gaussian, linear, and projection kernels when the risk factor is categorical (top row) or continuous (bottom row). We also considered an implementation that maximized power by combining all three kernels (combined). At each pathway size (columns), there were 20 simulated data sets with 1,000 pathways and the empirical false positive rate was assessed at a type I error rate of 0.05 (red line).

**Figure S3:**
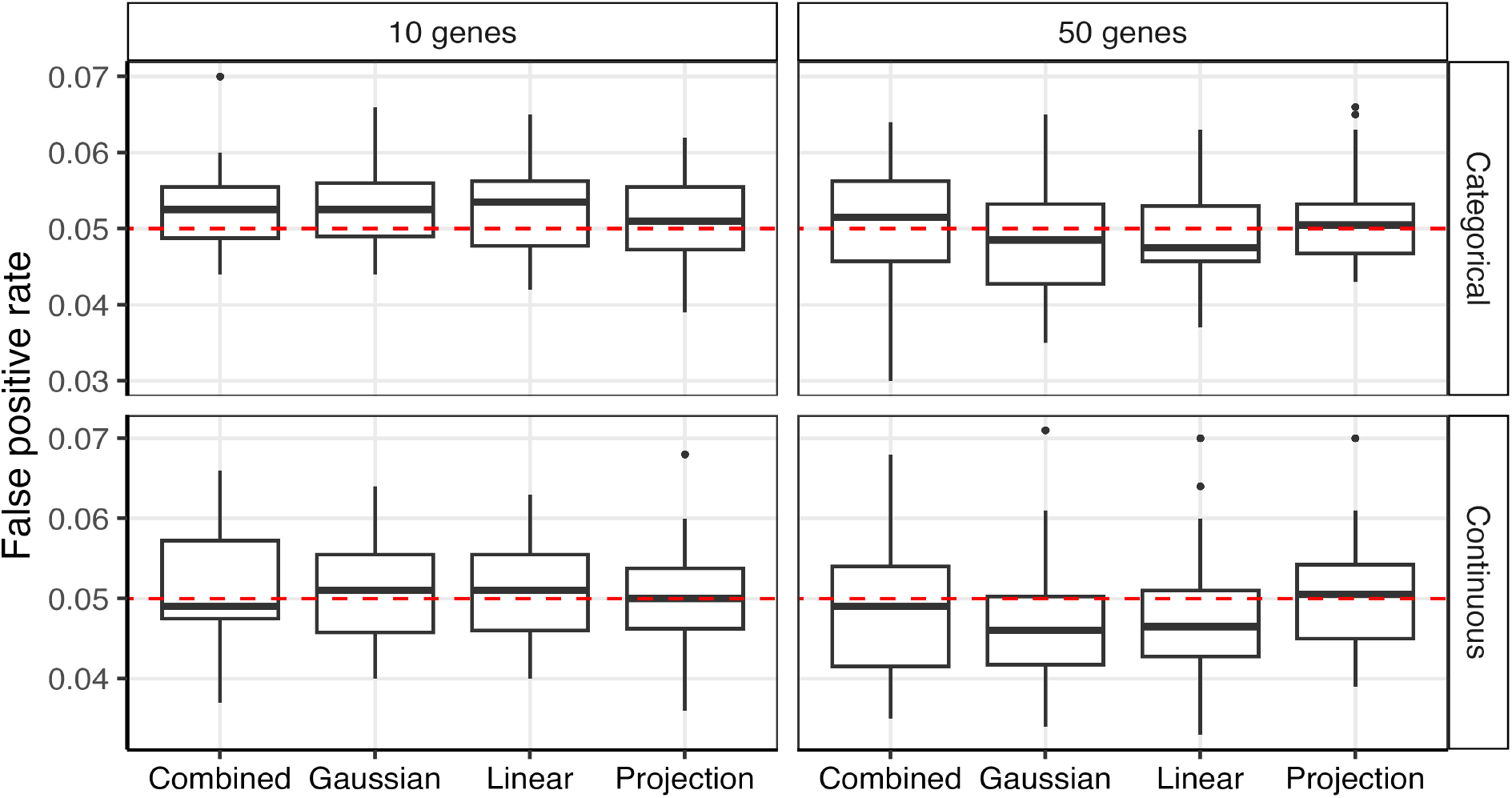
Assessing type I error rate in the RNA-seq simulation setting. We applied KDCA using the Gaussian, linear, and projection kernels to categorical (top row) and continuous (bottom row) risk factors. We also considered an implementation that maximized power by combining all three kernels (combined). At each pathway size (columns), there were 20 simulated data sets with 1,000 pathways and the empirical false positive rate was assessed at a type I error rate of 0.05 (red line).

**Figure S4:**
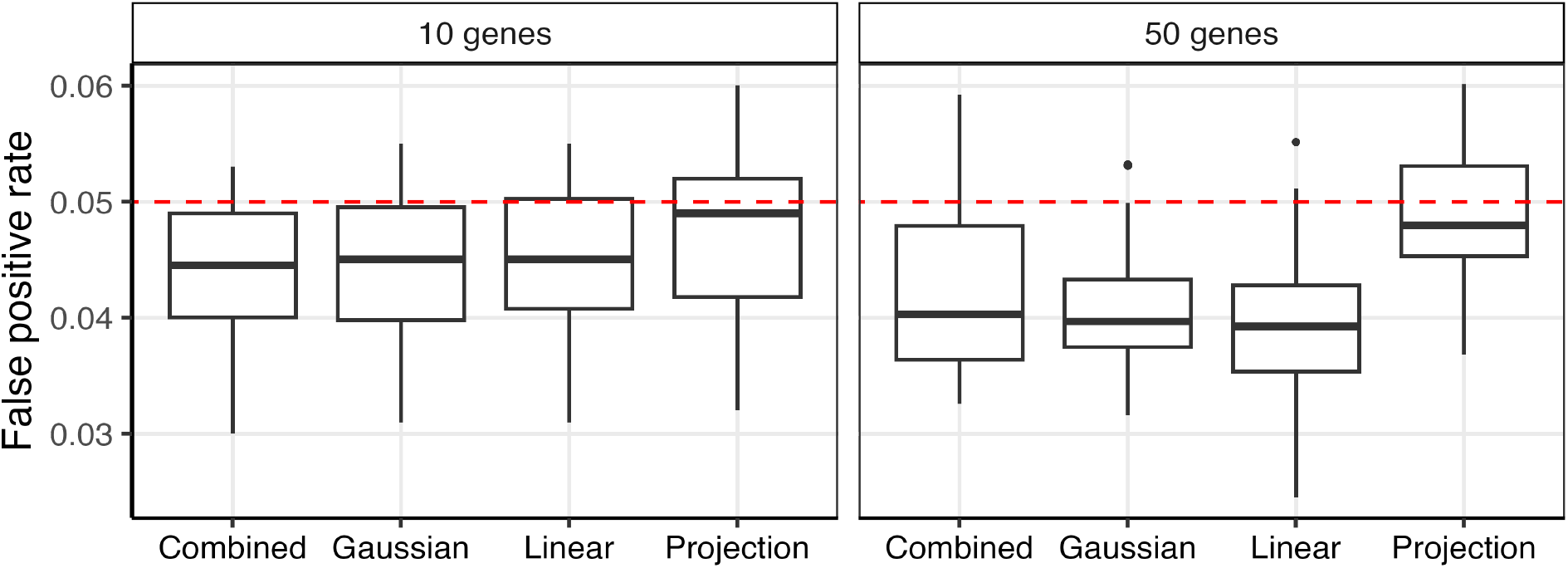
Assessing type I error rate when testing three risk factors for differential co-expression. We applied KDCA using the Gaussian, linear, and projection kernels to three primary variables. We also considered an implementation that maximized power by combining all three kernels (combined). At each pathway size (columns), there were 20 simulated data sets with 1,000 pathways and the empirical false positive rate was assessed at a type I error rate of 0.05 (red line).

**Figure S5:**
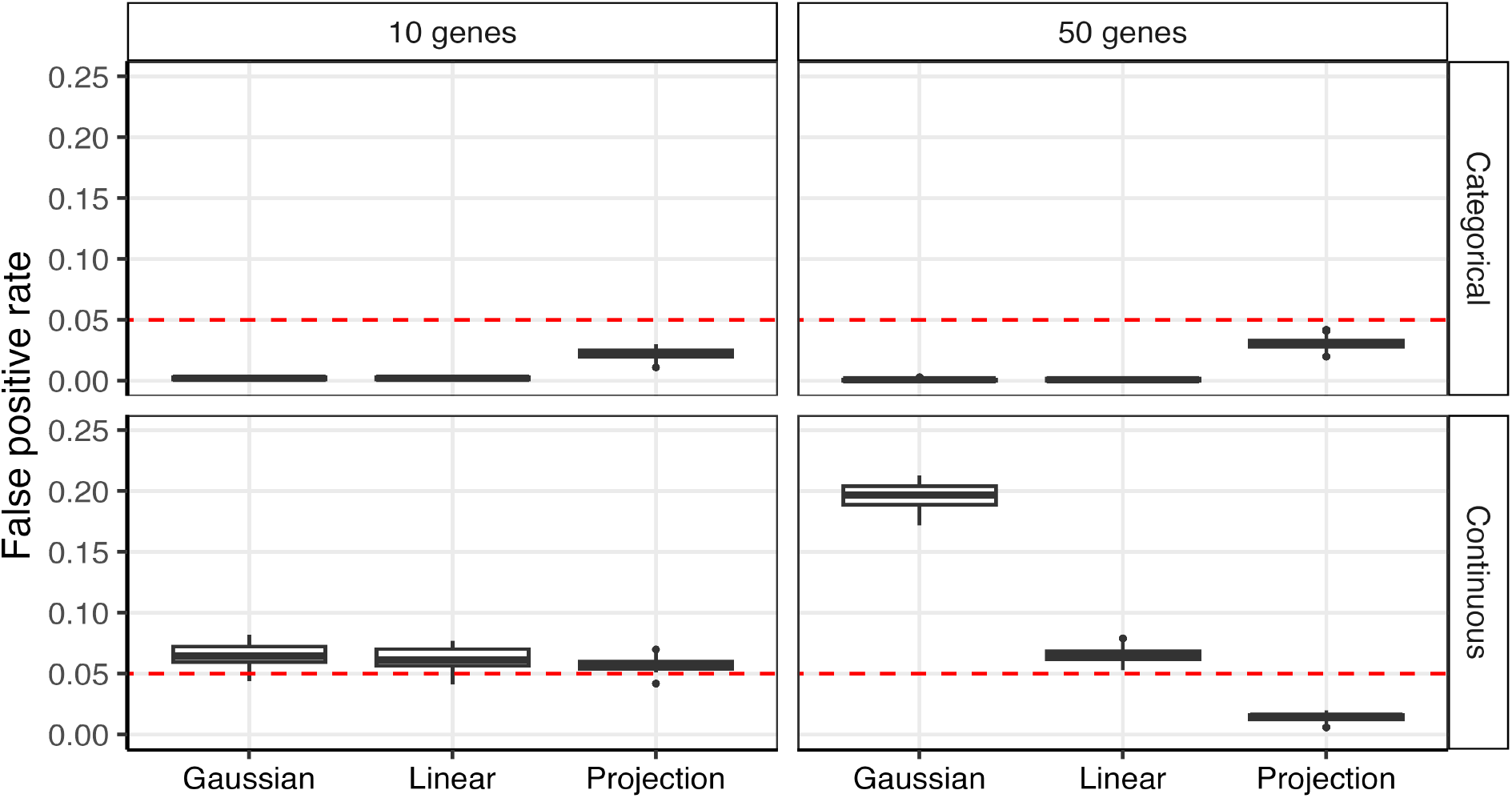
False positive rate of KDCA using a theoretical null approximation for the Gaussian, linear, and projection kernels when the risk factor is categorical (top row) or continuous (bottom row). At each pathway size (columns), there were 20 simulated data sets with 1,000 pathways and the empirical false positive rate was assessed at a type I error rate of 0.05 (red line).

**Figure S6:**
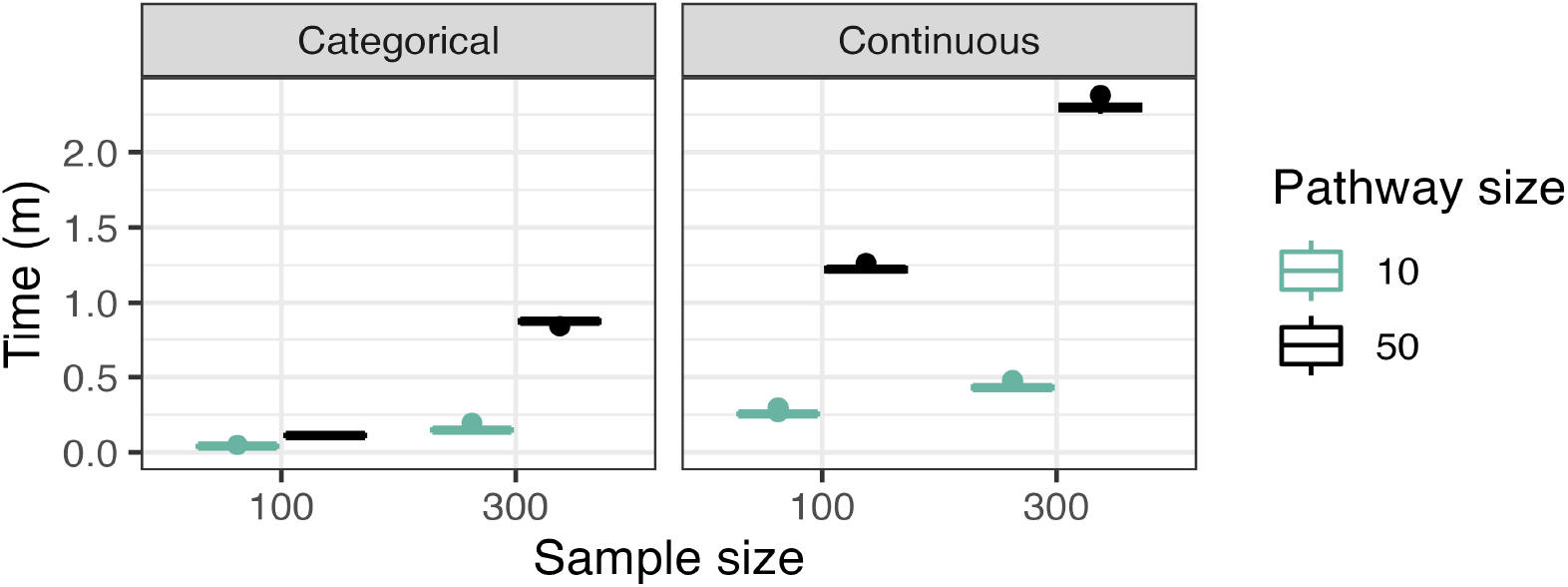
The computational time of KDCA with 1,000 permutations as a function of the sample size (x-axis), pathway size (color), and risk factor type (columns) in the simulation study (normal setting). There were 50 replicates at each setting and the simulations were performed on a single core of an Apple M3 processor.

**Figure S7:**
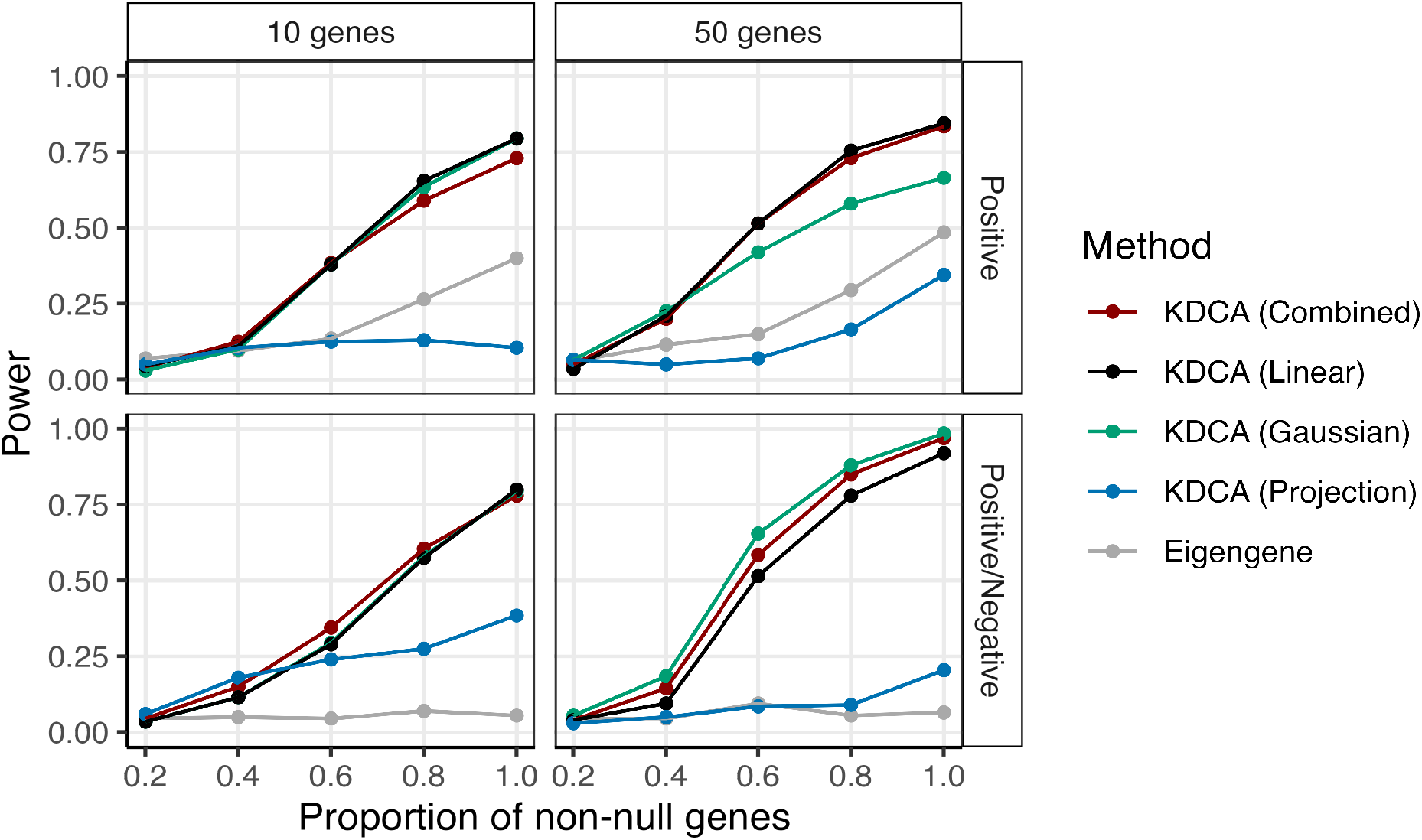
The empirical power of KDCA using the Gaussian (green), linear (black), and projection (blue) kernels when there are three risk factors. We also compared an implementation of KDCA that maximized power across all kernels (combined; red) to a standard eigengene (grey) approach. Our simulation study varied the pathway size (columns) and the type of differential co-expression (rows). Each point is the empirical power from 200 simulations at a significance threshold of 0.05.

**Figure S8:**
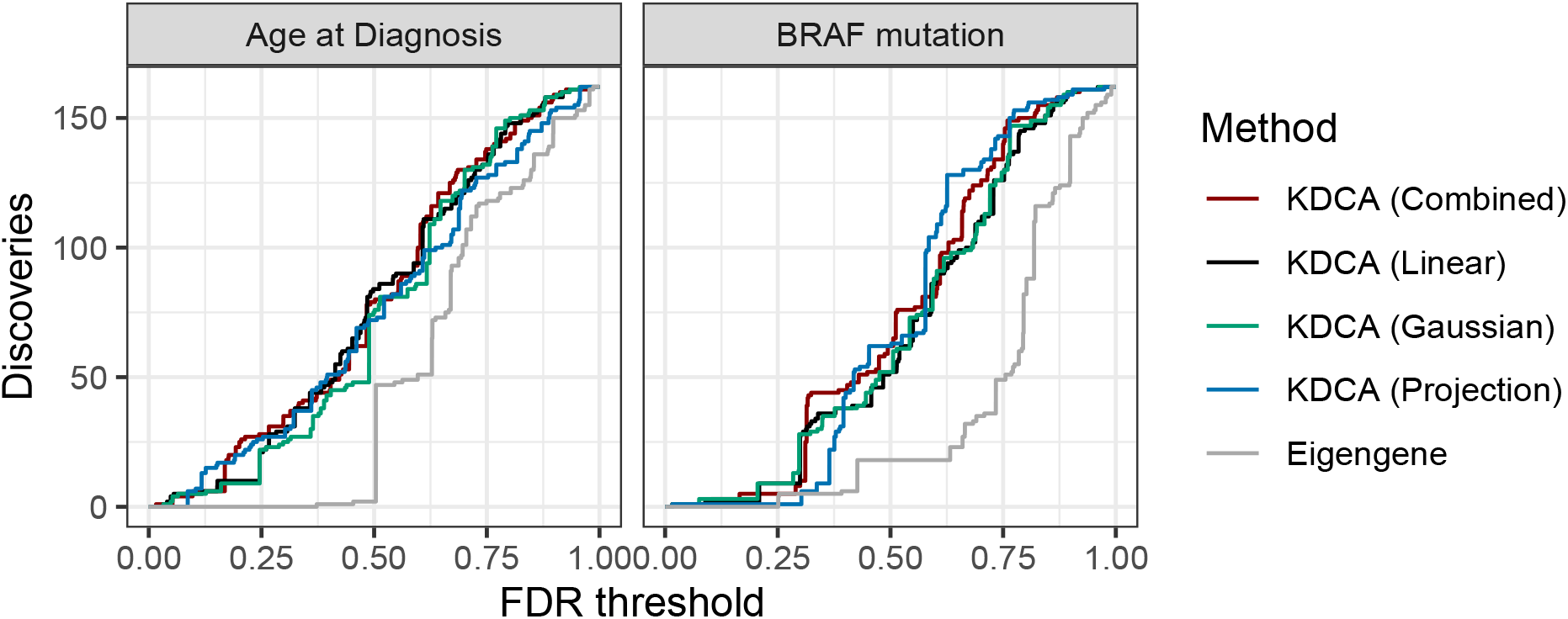
The number of discoveries as a function of FDR threshold using the Gaussian (green), linear (black), and projection (blue) kernels in the TCGA thyroid cancer data set using the BioCarta pathways. We implemented an aggregate version (red) to maximize power and compared KDCA to an eigengene approach (grey). We tested *BRAF* mutation status (right) and age of diagnosis (left) for differential co-expression.

### 5.3 Supplementary tables

**Table S1:**
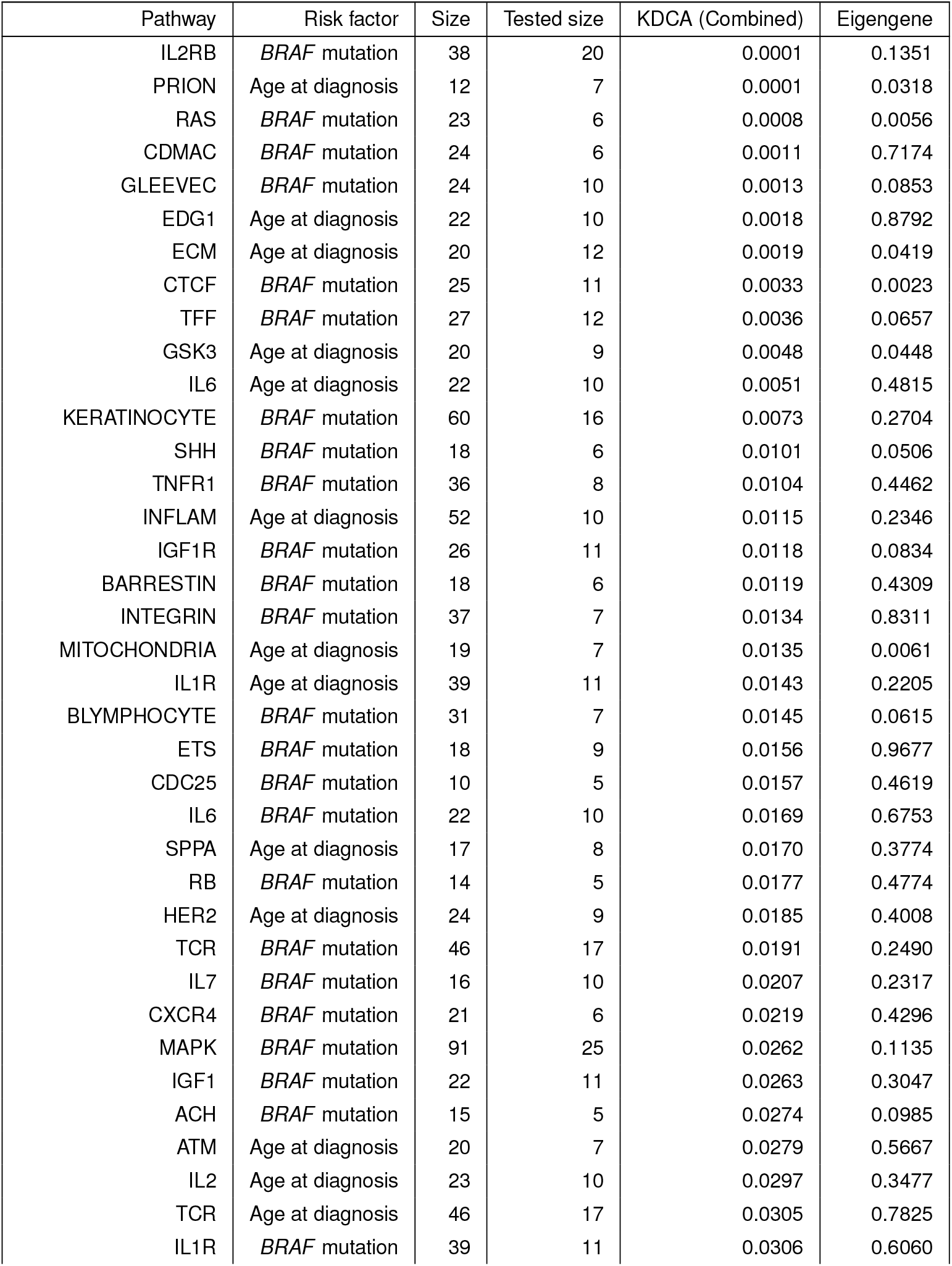
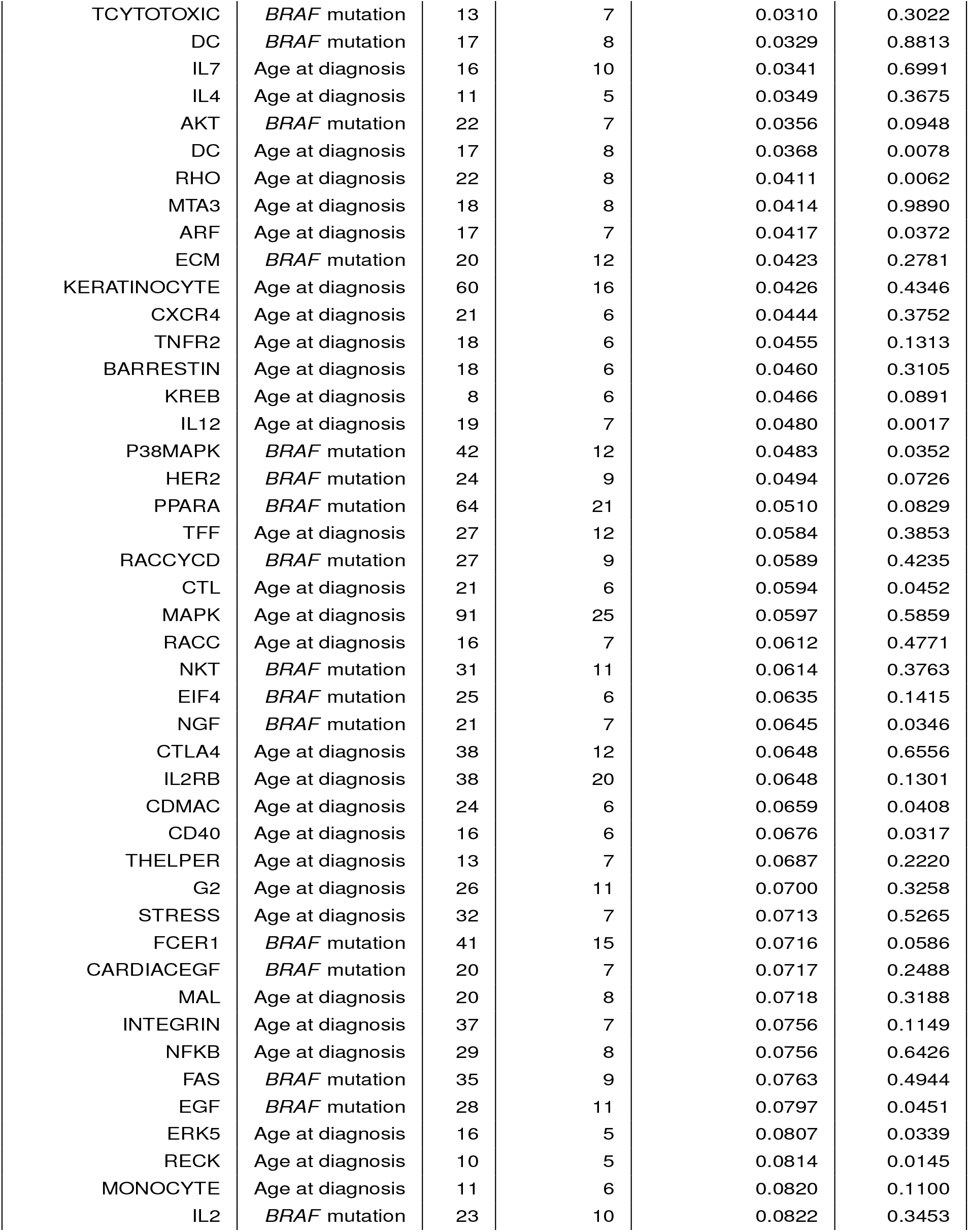
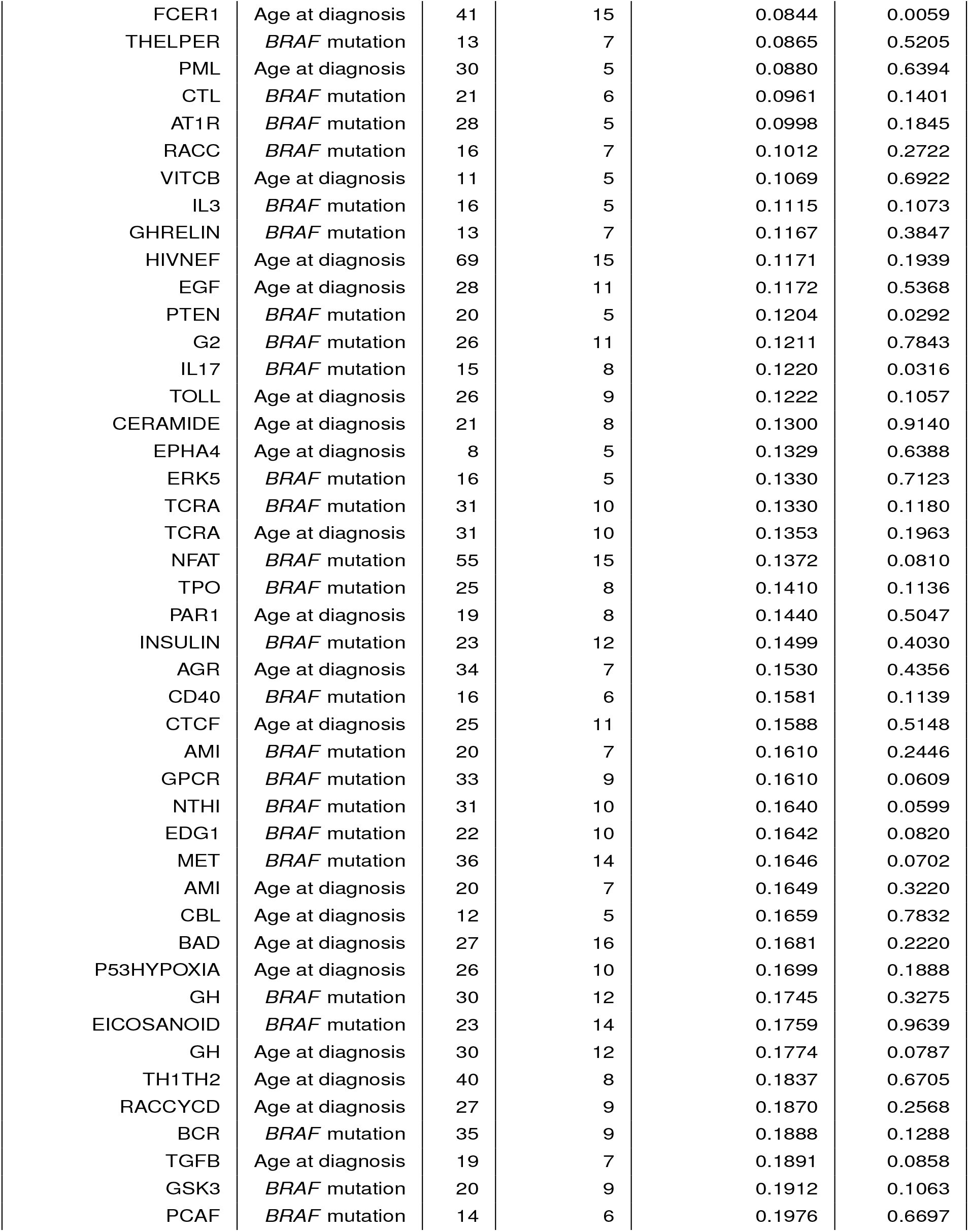
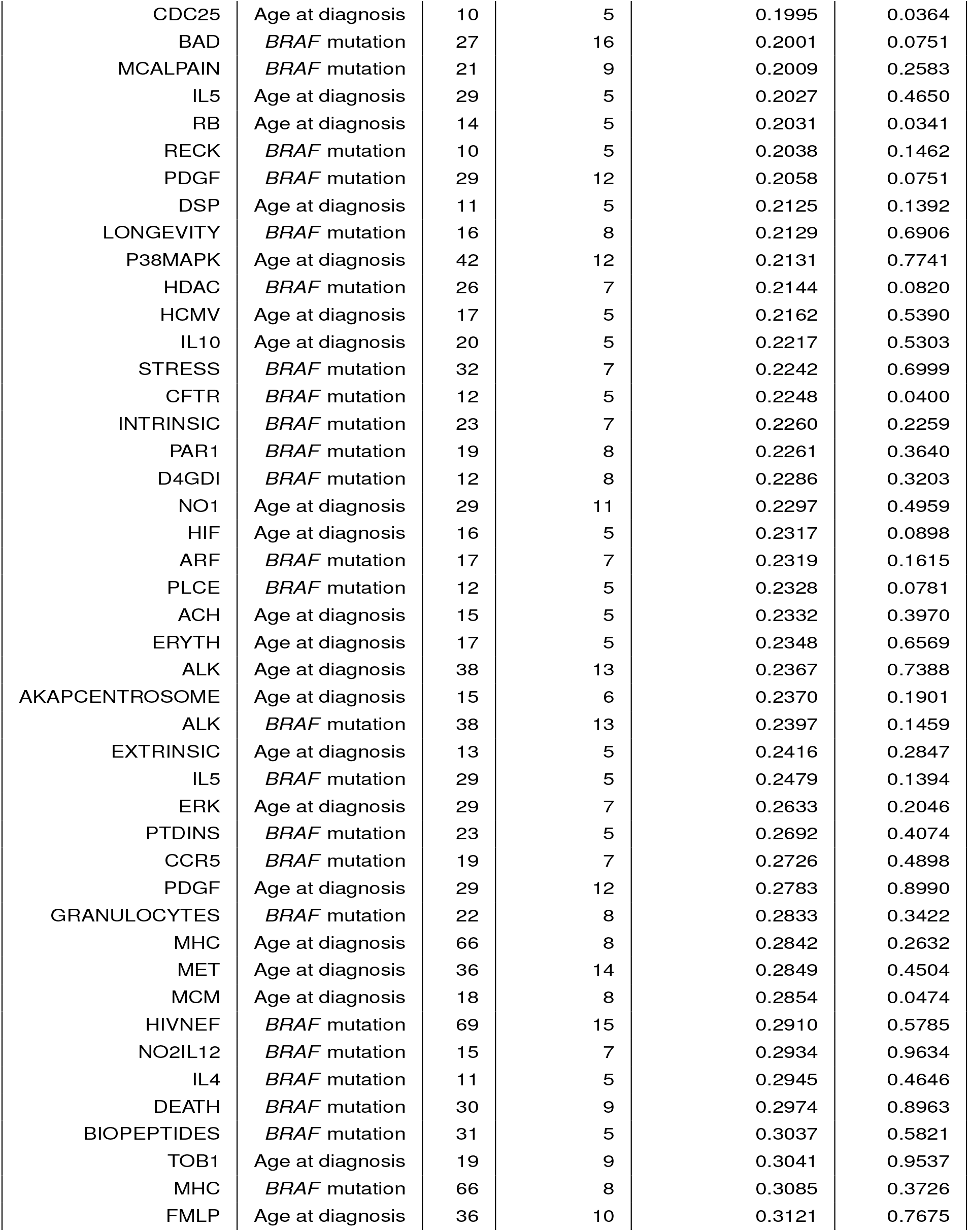
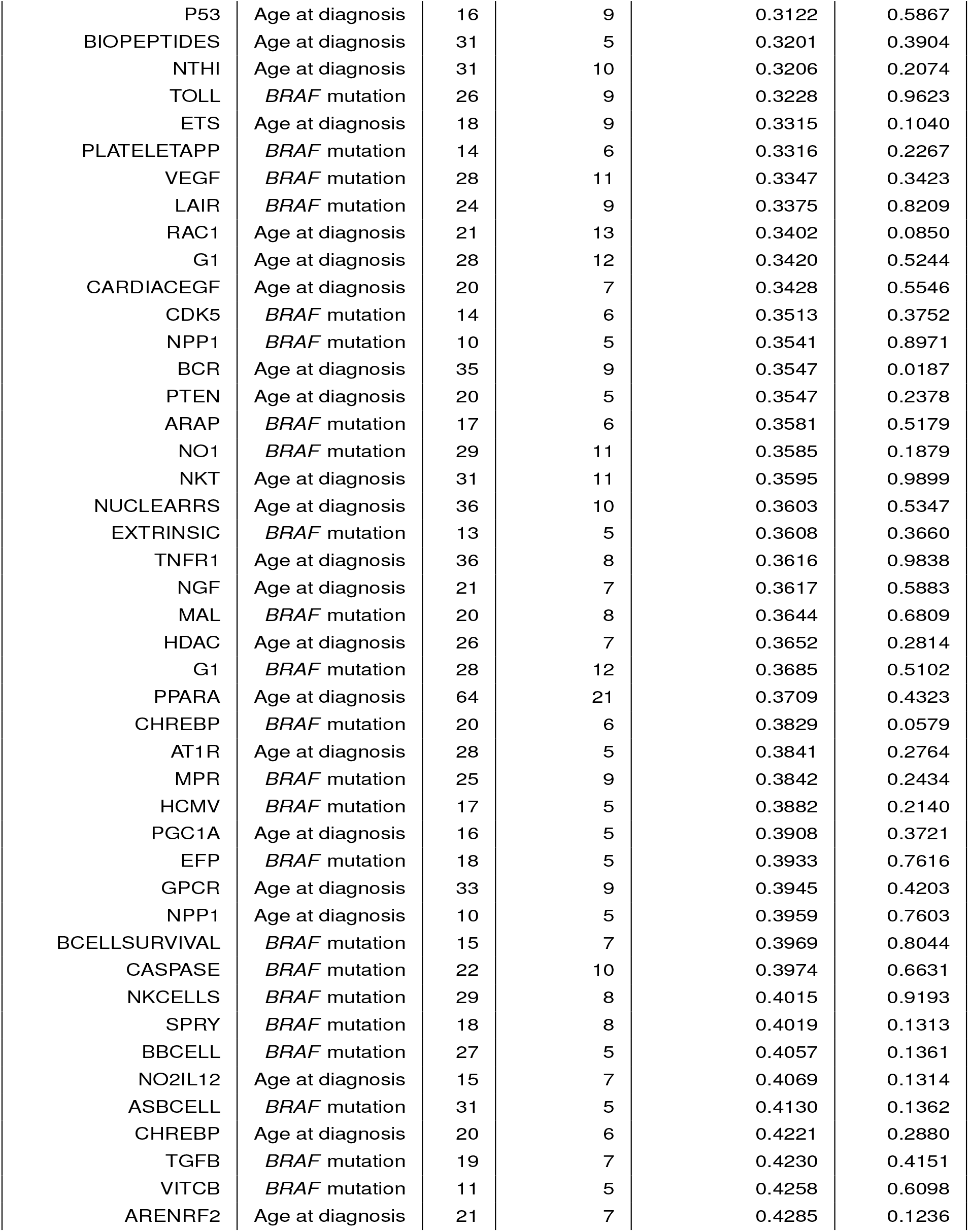
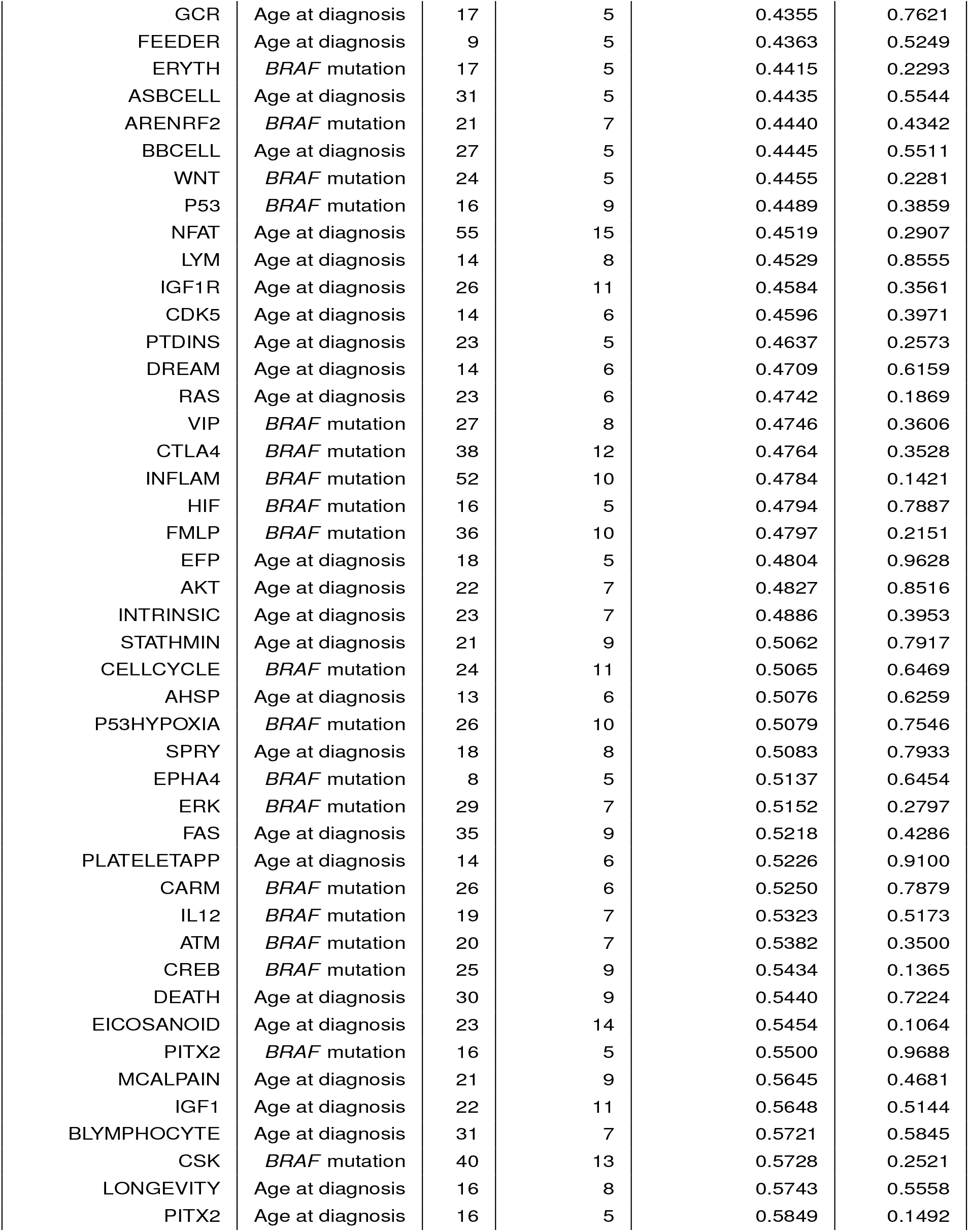
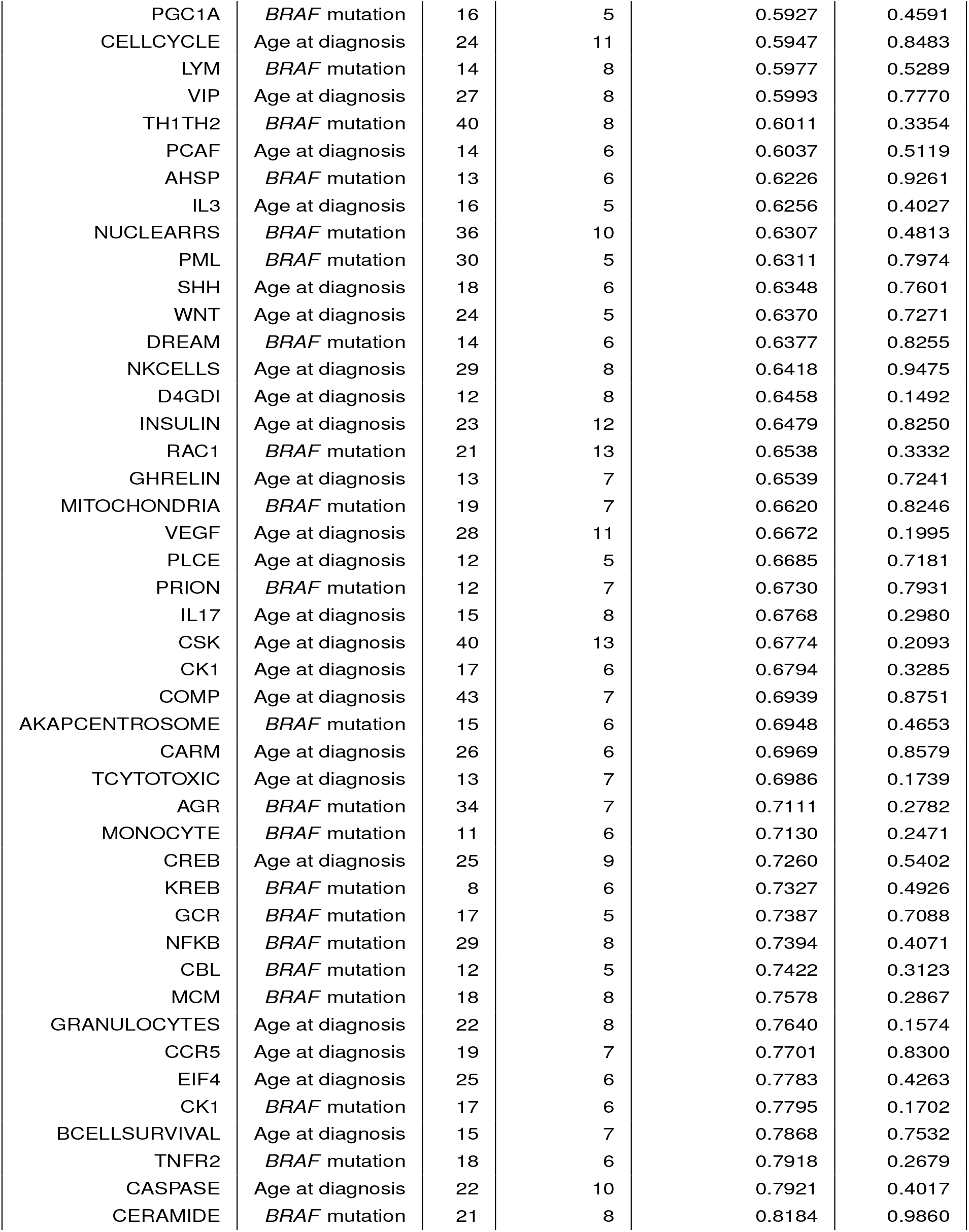
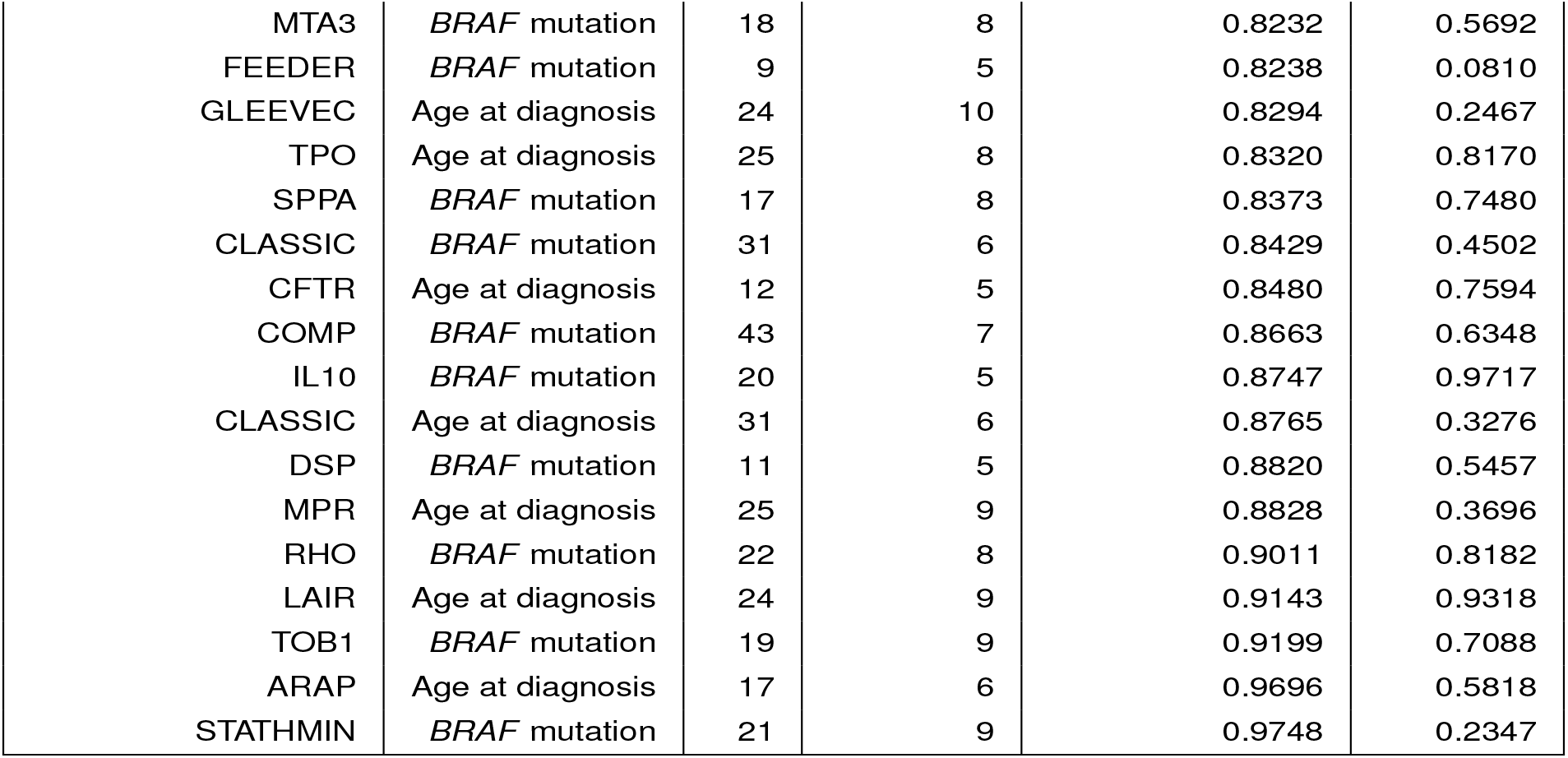
Significance results of the TCGA thyroid cancer data using age of diagnosis and *BRAF* mutation as the risk factors of interest. The *p*-values for KDCA (Combined) and an eigengene approach are reported for each pathway tested.

## Notes

### Competing Interest Statement

The authors have declared no competing interest.

